# Concerted activities of Mcm4, Sld3 and Dbf4 in control of origin activation and DNA replication fork progression

**DOI:** 10.1101/021832

**Authors:** Yi-Jun Sheu, Justin B. Kinney, Bruce Stillman

## Abstract

Eukaryotic chromosomes initiate DNA synthesis from multiple replication origins in a temporally specific manner during S phase. The replicative helicase Mcm2-7 functions in both initiation and fork progression and thus is an important target of regulation. Mcm4, a helicase subunit, possesses an unstructured regulatory domain that mediates control from multiple kinase signaling pathways, including the Dbf4-dependent Cdc7 kinase (DDK). Following replication stress in S phase, Dbf4 and Sld3, an initiation factor and essential target of Cyclin-Dependent Kinase (CDK), are targets of the checkpoint kinase Rad53 for inhibition of initiation from origins that have yet to be activated, so-called late origins. Here, whole genome DNA replication profile analysis is employed to access under various conditions the effect of mutations that alter the Mcm4 helicase regulatory domain and the Rad53 targets, Sld3 and Dbf4. Late origin firing occurs under genotoxic stress when the controls on Mcm4, Sld3 and Dbf4 are simultaneously eliminated. The regulatory domain of Mcm4 plays an important role in the timing of late origin firing, both in an unperturbed S phase and dNTP limitation. Furthermore, checkpoint control of Sld3 impacts fork progression under replication stress. This effect is parallel to the role of the Mcm4 regulatory domain in monitoring fork progression. Hypomorph mutations in *sld3* are suppressed by a *mcm4* regulatory domain mutation. Thus, in response cellular conditions, the functions executed by Sld3, Dbf4 and the regulatory domain of Mcm4 intersect to control origin firing and replication fork progression, thereby ensuring genome stability.

## INTRODUCTION

Eukaryotic cells initiate DNA synthesis from multiple replication origins on each chromosome to ensure efficient duplication of the genome in S phase. Activation of replication origins is achieved through two distinct steps that take place at separate stages of the cell division cycle. The first step, licensing of replication origins, occurs in G1 when CDK activity is low (Diffley 2011). During this process, a double hexameric minichromosome maintenance (MCM) complex, composed of two Mcm2-7 hexamers, is loaded onto each replication origin to form a pre-Replicative Complex (pre-RC) by the Origin Recognition Complex (ORC) and licensing factors, Cdc6 and Cdt1 (Diffley 2011). The second step, activation of licensed origins, occurs at each origin in a temporally controlled manner throughout S phase and requires activities of two S phase kinases, the S phase Cyclin-dependent Kinases (CDKs) and the Dbf4-dependent Cdc7 kinase (DDK) (Tanaka and Araki 2013). CDK phosphorylates two key substrates, Sld2 and Sld3, and promotes their binding to Dpb11 (Tanaka et al. 2007; Zegerman and Diffley 2007). DDK phosphorylates several subunits of the Mcm2-7 hexamer and, most importantly, blocks an intrinsic inhibitory activity residing within the amino-terminus of the Mcm4 subunit (Sheu and Stillman 2006; Randell et al. 2010; Sheu and Stillman 2010). The action of these S phase kinases facilitates recruitment of Cdc45 and the GINS complex, composed of protein subunits Sld5, Psf1, Psf2 and Psf3, to the inactive MCM double hexamer and converts it into an active helicase complex, composed of Cdc45, Mcm2-7 and GINS (the CMG complex) (Tanaka and Araki 2013). The two-step process separates the loading and activation of replicative helicases at origins and thereby ensures that initiation from each origin occurs once and only once during each cell division cycle. Once origins are fully activated, the double helix unwinds and DNA polymerase and other replisome components are recruited to establish replication forks, where new DNA is copied bi-directionally from each origin.

Initiation of DNA synthesis from licensed origins across the genome (origin firing) follows a pre-determined temporal pattern (Rhind and Gilbert 2013). In budding yeast, the timing of DNA replication can be traced to the activation of individual origins. Origin activation occurs continuously during S phase, but those that fire first in S phase are referred to as early origins and those that fire later are late origins. Despite being an essential target of CDK, Sld3, together with Sld7 and Cdc45, binds to the loaded Mcm2-7 hexamer in a manner dependent on DDK but not CDK (Heller et al. 2011; Tanaka et al. 2011). This association is a prerequisite for the subsequent CDK-dependent recruitment of a pre-loading complex, composed of Sld2, Dpb11, GINS and pol ε (Muramatsu et al. 2010). It was proposed that DDK-dependent recruitment of the limiting Sld3-Sld7-Cdc45 is a key step for determining the timing of origin firing (Tanaka et al. 2011). Furthermore, simultaneous overexpression of several limiting replication factors advances late origin firing (Mantiero et al. 2011; Tanaka et al. 2011).

Under genotoxic stress during S phase, DNA damage checkpoint pathways inhibit late origin firing (Zegerman and Diffley 2009). In budding yeast, DNA damage activates the mammalian ATM/ATR homolog, Mec1 kinase, which in turn activates the Rad53 effector kinase (the homolog of mammalian Chk2) to phosphorylate and inhibit the activities of Sld3 and Dbf4, thereby preventing late origin firing (Lopez-Mosqueda et al. 2010; Zegerman and Diffley 2010). Some firing of late origins could be detected under DNA damaging conditions in phosphorylation mutants of these two targets rendered refractory to the inhibition by Rad53. An initiation inhibitory activity within the non-structured, amino-terminal regulatory domain of Mcm4 (NSD; Fig. 1) also plays a role in regulating origin firing under genotoxic stress (Sheu et al. 2014). Because this domain is a target of DDK (Masai et al. 2006; Sheu and Stillman 2006; Sheu and Stillman 2010), it is conceivable that Mcm4 could mediate the checkpoint control by Rad53 phosphorylation of Dbf4. However, since DDK has targets other than Mcm4 and Mcm4 is regulated by signals in addition to DDK, a more comprehensive picture of how these factors cooperate to control origin firing under stress condition remains to be addressed.

**Figure 1.**
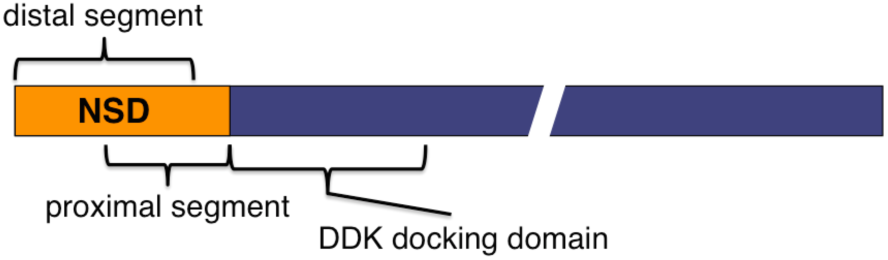
Diagram of the Mcm4 subunit of Mcm2-7 helicase. The two overlapping segments within the Mcm4 structurally disordered N-terminal serine/threonine-rich domain (NSD) are shown.

In addition to origin activation, DNA synthesis can be controlled at the level of replication fork progression. For example, deoxyribonucleoside triphosphate (dNTP) levels influence the rate of replication fork progression (Santocanale and Diffley 1998; Alvino et al. 2007). Hydroxyurea (HU) inhibits the activity of ribonucleotide reductase (RNR) and causes a dramatic slowdown of replication fork progression. In contrast, high dNTP concentration inhibits ORC-dependent initiation of DNA replication (Chabes and Stillman 2007). It has been proposed that dNTP levels are key determinants of replication fork speed and that cells adapt to replication stress by up-regulating dNTP pools (Poli et al. 2012). Methyl methanesulfonate (MMS), a DNA-alkylating agent, also results in slower fork progression while activating the DNA damage checkpoint response (Tercero and Diffley 2001). Although Mec1 and Rad53 are essential for preventing DNA replication fork catastrophe, these checkpoint kinases are not required for fork slowing in MMS (Tercero and Diffley 2001; Tercero et al. 2003). Thus, it is possible that an alternative mechanism might regulate fork progression under stress conditions.

The structurally disordered N-terminal serine/threonine-rich domain (NSD) of Mcm4 participates in both initiation and fork progression (Sheu et al. 2014). It can be subdivided into two overlapping but functionally distinct segments, the proximal segment and the distal segment (Fig. 1). The proximal segment of the NSD (amino acids 74-174) is responsible for the initiation inhibitory activity that is mitigated by DDK through phosphorylation (Sheu and Stillman 2006; Sheu and Stillman 2010). The distal segment (amino acids 2-145) is important for controlling fork progression and checkpoint response under replication stress caused by depletion of dNTP pools and its function is regulated by CDK (Devault et al. 2008; Sheu et al. 2014). Thus, this intrinsic regulatory domain of the replicative helicase may cooperate with additional factors to control origin firing and replication fork progression in response to various environmental conditions. Herein, we examine the contributions of Mcm4, Sld3 and Dbf4 in control of DNA damage induced control of both origin activation and DNA replication fork progression.

## RESULTS

### Eliminating controls on Mcm4, Sld3 and DDK is required for maximal late origin firing in hydroxyurea

To evaluate how deregulating the control on Mcm4, Sld3 and Dbf4 impact origin firing on a genome-wide scale, the replication profiles of the wild type (*WT*) and mutant strains with \*\*\**mcm4*^*Δ74-174*^ [*mcm4* mutant lacking the proximal NSD domain, the target of DDK], *sld3-38A* and *dbf4-19A* [alleles of *SLD3* and *DBF4*, respectively, that are resistant to the checkpoint control due to serine/threonine to alanine substitutions at the Rad53 target sites (Zegerman and Diffley 2010)], were analyzed in single, double and triple mutant combinations. Cells were analyzed by releasing them synchronously from G1 phase into S phase in the presence of HU for 90 min (Fig. 2, data for chromosome IV). Late origins were inactive at this time in *WT* (Fig. 2A, profile *WT;* red arrows) while a very low level of late origin firing was detected in each of the single mutants (profiles *mcm4*^*Δ74-174*^, *sld3-38A* or *dbf4-19A*; Fig. 2A), consistent with previous findings (Lopez-Mosqueda et al. 2010; Zegerman and Diffley 2010; Sheu et al. 2014). Late origin firing appeared more prominent in all of the double mutant combinations (profiles *mcm4Δ*^*74-174*^ *sld3-38A*, *sld3-38A dbf4-19A* and *mcm4Δ*^*74-174*^ *dbf4-19A*), suggesting that these three factors function in pathways that are not completely overlapping. Among the double mutants, the *mcm4Δ*^*74-174*^ *sld3-38A* combination showed the most robust late origin firing. Thus it is likely that Mcm4 and Sld3 function in separate control pathways to regulate origin firing. The *mcm4*^*Δ74-174*^ *dbf4-19A* combination only increased late origin firing slightly compared with each single mutant, consistent with the finding that Dbf4 and Mcm4 act in pathways that overlap extensively as Mcm4 is the essential target of DDK (Sheu and Stillman 2010). The detectable, but limited increase in late origin firing in *sld3-38A dbf4-19A* cells in comparison with their single mutants also suggests overlapping of the pathways involving Sld3 and Dbf4. In the triple mutant (Fig. 2A profile *mcm4*^*Δ74-174*^ *sld3-38A dbf4-19A*), many late origins fire very robustly, more than any of the double and single mutants and the efficiency of late origin firing approached the level of the early origins. The massive firing of late origins in the triple mutant further suggests that each of the three factors contribute independently to control of origin firing through overlapping but non-identical pathways.

**Figure 2.**
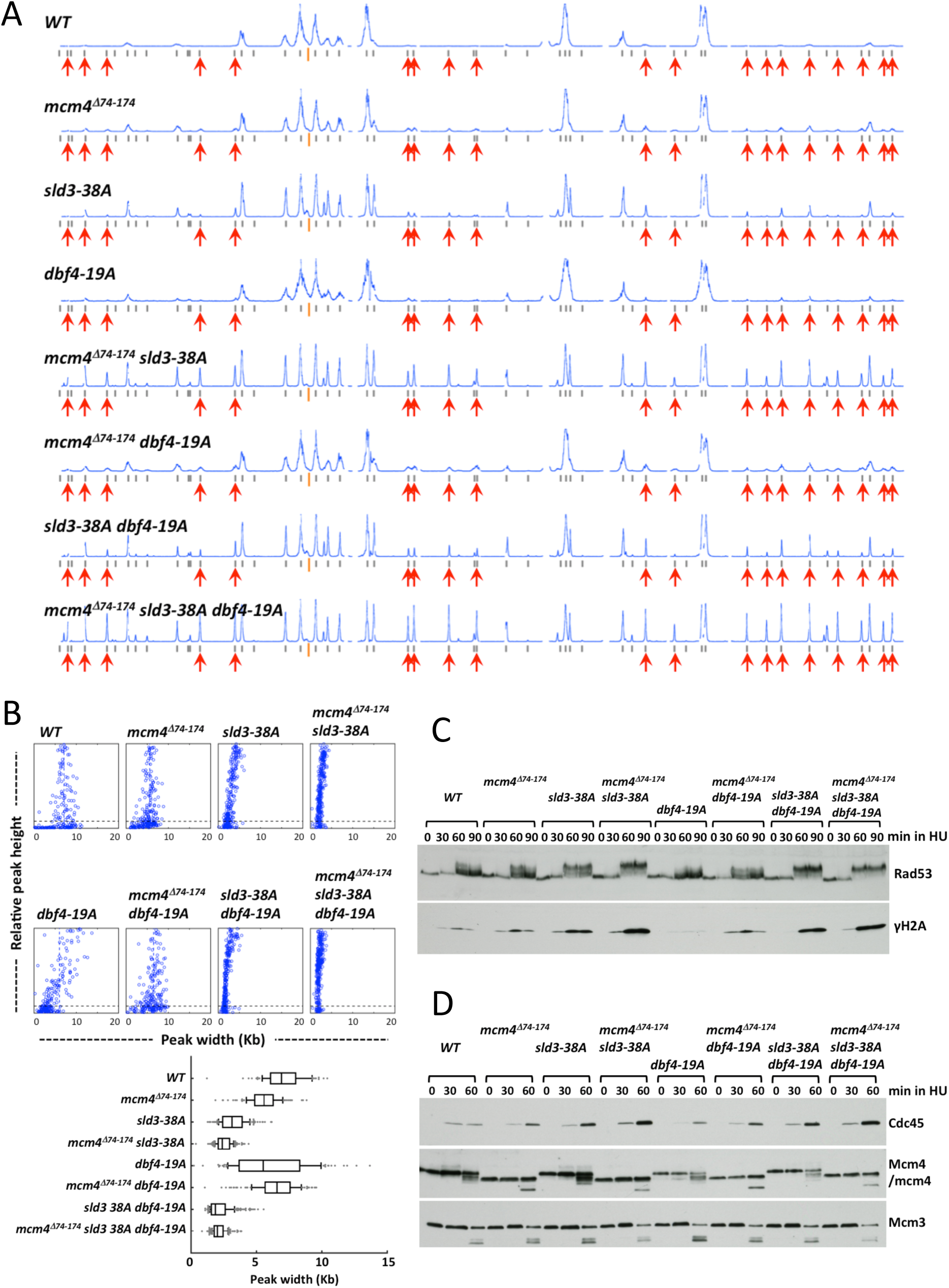
Simultaneously alleviating controls on Mcm4, Sld3 and Dbf4 allows efficient late origin firing in HU despite an elevated checkpoint response. (A and B) Yeast cells were synchronized in G1 phase and released in to YPD containing 0.2 M HU and 0.5 mM EdU for 90 min. (A) Replication profiles of chromosome IV for the wild type (*WT*), single, double and triple mutants with *mcm4*^*Δ74-174*^, *sld3 38A* and/or *dbf4 19A* alleles as indicated on the top-left of each profile. Gray bars under each profile indicate annotated replication origins according to the OriDB (full name and REF). Orange bars indicate the position of the centromere. Red arrows point out some late origins that are inactive in the wild type cells but fire in the mutants, in particularly the triple mutant. (B) Distribution of fork progression from activated origins. Top panels show the peak width-height plots of all the recorded origins across the entire genome for each strain in (A). Box plot shows the fork progression, excluding peaks with heights smaller than 10 % of the maximal height (also shown as black dash line in individual width-height plots). Box and whiskers indicate 25–75 and 10–90 percentiles, respectively. (C) Cells of the indicated yeast strains were synchronized in G1, released into 0.2 M HU and collected at the indicated time points. Protein samples were prepared using TCA extraction and analyzed by immunoblot. (D) Analysis of chromatin-bound proteins. Cells synchronized in G1 were released into 0.2 M HU and collected at the indicated time points. Chromatin-bound proteins were extracted from cells and analyzed by immunoblot.

Late origin firing in these mutants was not due to defects in the HU induced checkpoint response. Judging from levels of Rad53 hyper-phosphorylation and phosphorylation of S129 in histone H2A (γH2A), the checkpoint signaling in *mcm4*^*Δ74-174*^, *sld3-38A* and all the double and triple mutants was stronger than *WT* (Fig. 2C, Rad53 and γH2A). The elevated checkpoint signaling could be also reflect more origin firing in these cells and more stalled forks (Tercero 2003). Consistent with this idea, higher levels of Cdc45 loading, but not Mcm3 levels were also detected on S phase chromatin in these mutants (Fig. 2D).

### Proximal segment of Mcm4 NSD delays late origin firing in HU when control of Sld3 and Dbf4 by checkpoint is abrogated

To determine whether the triple mutant activated late origins with the same kinetics as early origins, the replication profiles of *WT* and the triple mutant were compared at different time points after release from G1 phase into S phase in the presence of HU (Fig. 3). At 25 min after release into HU, early origins fired in *WT* cells, while late origins remained inactive (Fig. 3A). At 50 min, the profile of activated origins in *WT* remained similar to that at 25 min, with a small increase of peak width, indicating progression of replication forks. At 75 min, replication forks progressed further, but the pattern of origin firing remained unchanged. In contrast, the triple mutant activated some late origins by 25 min. and the peak height of late origins continued to increase relative to that of early origins as time progressed to 50 and 75 min, suggesting that late origins continued to fire in this cell population. These data show that removing the proximal NSD segment of Mcm4, together with abolishing Rad53 phosphorylation of Sld3 and Dbf4 allowed late origins to fire efficiently in the presence of HU, but still in a temporally specific manner. In the presence of the proximal NSD segment, however, only low levels of late origin firing were detected in the *sld3-38A dbf4-19A* mutant at 50 min after release from G1 and only slightly increased at 75 min., but did not reach the same level as the *mcm4*^*Δ74-174*^ *sld3-38A dbf4-19A* mutant (Fig. 3B). These observations suggest that the proximal segment of the Mcm4 NSD prevents late origin firing at the earlier time in S phase despite the absence of active checkpoint inhibition of Sld3 and DDK function.

**Figure 3.**
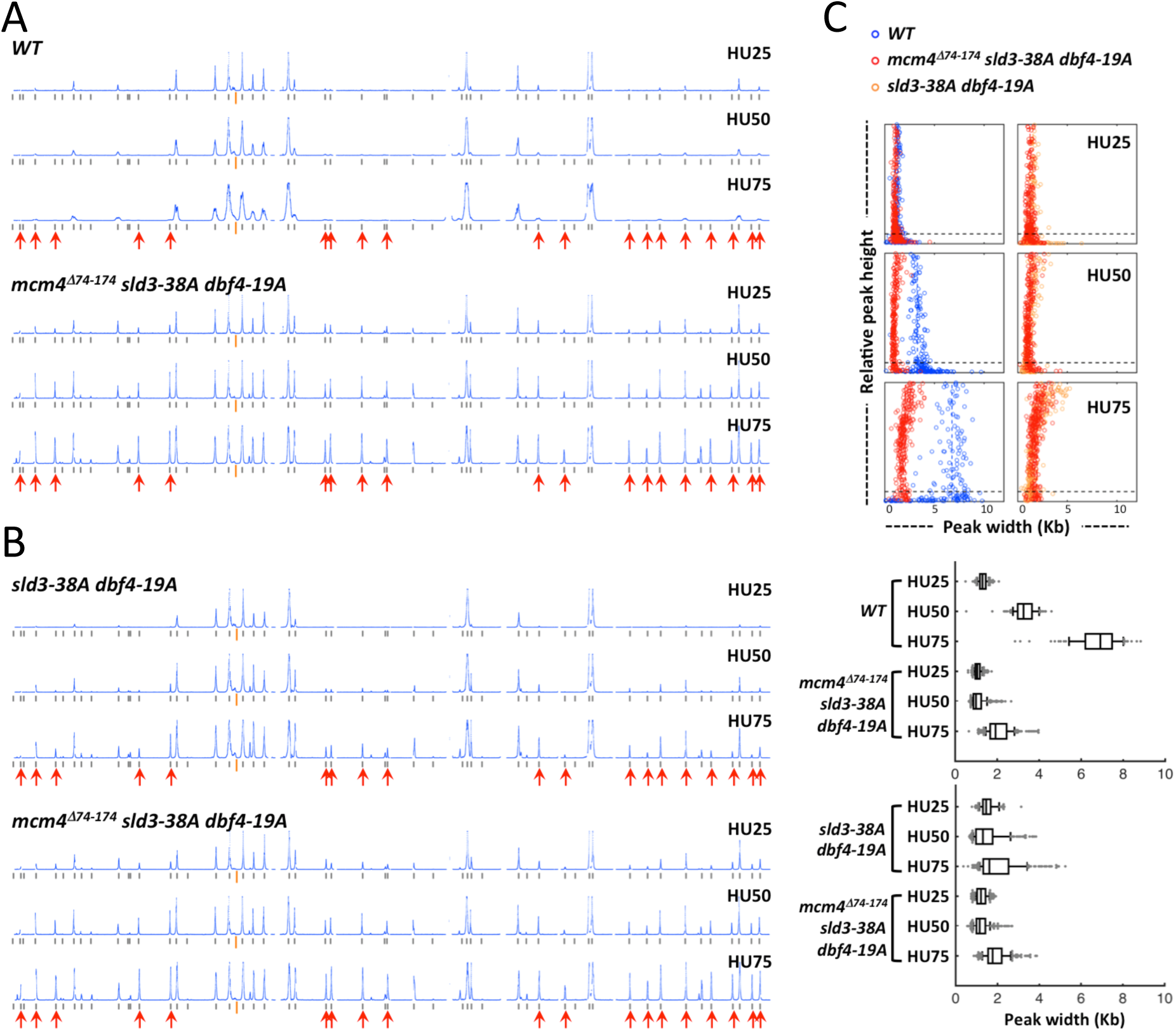
Proximal segment of the Mcm4 NSD delays late origin firing in HU when checkpoint control on Sld3 and Dbf4 is relieved. Yeast cells were synchronized in G1 phase and released in to YPD containing 0.2 M HU and 0.5 mM EdU. (A) Replication profiles of chromosome IV for *WT* and the *mcm4*^*Δ74-174*^ *sld3-38A dbf4-19A* triple mutant at 25, 50 and 75 min after release into HU from G1 arrest. (B) Replication profiles of chromosome IV for the *sld3-38A dbf4-19A* and *mcm4*^*Δ74-174*^ *sld3-38A dbf4-19A* mutants at 25, 50 and 75 min after release into HU from G1. (C) Distribution of fork progression from activated origins at the indicated time point for the wild type (blue, data from 3A), *sld3-38A dbf4-19A* (tan, data from 3B) and *mcm4*^*Δ74-174*^ *sld3-38A dbf4-19A* (red; left panels, data from 3A; right panels, data from 3B) mutants. Box graph shows the fork progression, excluding peaks with heights smaller than 10 % of the maximal height scale.

### The distal segment of Mcm4 NSD and Rad53 phosphorylation of Sld3 affect replication fork progression

In addition to revealing patterns of origin activation, whole genome replication profile analysis also provides information on the average replication fork progression from each origin in the population of cells. For computational analysis, we defined fork progression as the observed peak width at the half maximum of the peak height for each origin in the profile (Sheu et al. 2014). Analysis of replication profiles in HU showed that replication fork progression was much less in all the mutants containing the *sld3-38A* allele than in the wild type (Fig. 2A, 2B, 3C, 4A and 4B; Supplementary table 1), suggesting that phosphorylation of Sld3 by the checkpoint kinase Rad53 is needed to allow replication fork progression in HU. The *mcm4*^*Δ74-174*^ mutant lacking the Mcm4 proximal NSD domain also had less fork progression than the wild type, but the difference was subtle, yet reproducible (Fig. 2B and 4B; Supplementary table 1). Fork progression in the other Rad53 target mutant *dbf4-19A* appeared heterogeneous (Fig. 2B). Double mutants *sld3-38A dbf4-19A* and *mcm4*^*Δ74-174*^ *dbf4-19A* exhibited fork progression patterns resembling single mutants *sld3-38A* and *mcm4*^*Δ74-174*^, respectively, rather than that of the *dbf4-19A* mutant, suggesting that the functions of *sld3-38A* and *mcm4*^*Δ74-174*^ in fork progression are epistatic to that of *dbf4-19A*. Thus, Sld3 and the proximal NSD of Mcm4 are likely to function downstream of Dbf4 in regulation fork progression in HU.

Analysis of fork progression of samples from the time course experiment showed that the extent of progression at 25 min after release to HU was similar among *WT*, *sld3-38A dbf4-19A* and *mcm4*^*Δ74-174*^ *sld3-38A dbf4-19A* mutants (Fig. 3C; Supplementary table 1). At 50 min, replication forks advanced further in the wild type. However, in the *sld3-38A dbf4-19A* and *mcm4*^*Δ74-174*^ *sld3-38A dbf4-19A* mutants, the progression remained at the same level as those at 25 min. Replication fork progression continued in the wild type at 75 min while the forks in the mutant progressed very little.

In contrast to *sld3-38A* and *mcm4*^*Δ74-174*^, mutations affecting the function of the distal NSD of Mcm4, such as *mcm4*^*Δ2-145*^ and *mcm4*^*Δ74-174, 4(SP→AP)*^, exhibited extensive fork progression (Sheu et al. 2014) (see Fig. 4B). To establish the genetic relationship between the function of Sld3 and the distal segment of the NSD of Mcm4 in controlling replication fork progression, replication profiles in HU were analyzed in single and double mutants (Fig. 4). In the *mcm4*^*Δ2-145*^ *sld3-38A* double mutant, the opposing effects of the individual mutations (*mcm4*^*Δ2-145*^ and *sld3-38A*) appeared to cancel each other (Fig. 4A and B; Supplementary table S1), suggesting their controls operate through separate pathways. The *mcm4*^*Δ74-174, 4(SP→AP)*^ has mutations in the Mcm4 distal NSD segment and inactivates CDK phosphorylation sites (Sheu et al. 2014). The *mcm4*^*Δ74-174, 4(SP→AP)*^ *sld3-38A* double mutant and the *mcm4*^*Δ2-145*^ *sld3-38A* mutant showed a similar fork progression phenotypes (Fig. 4B).

**Figure 4.**
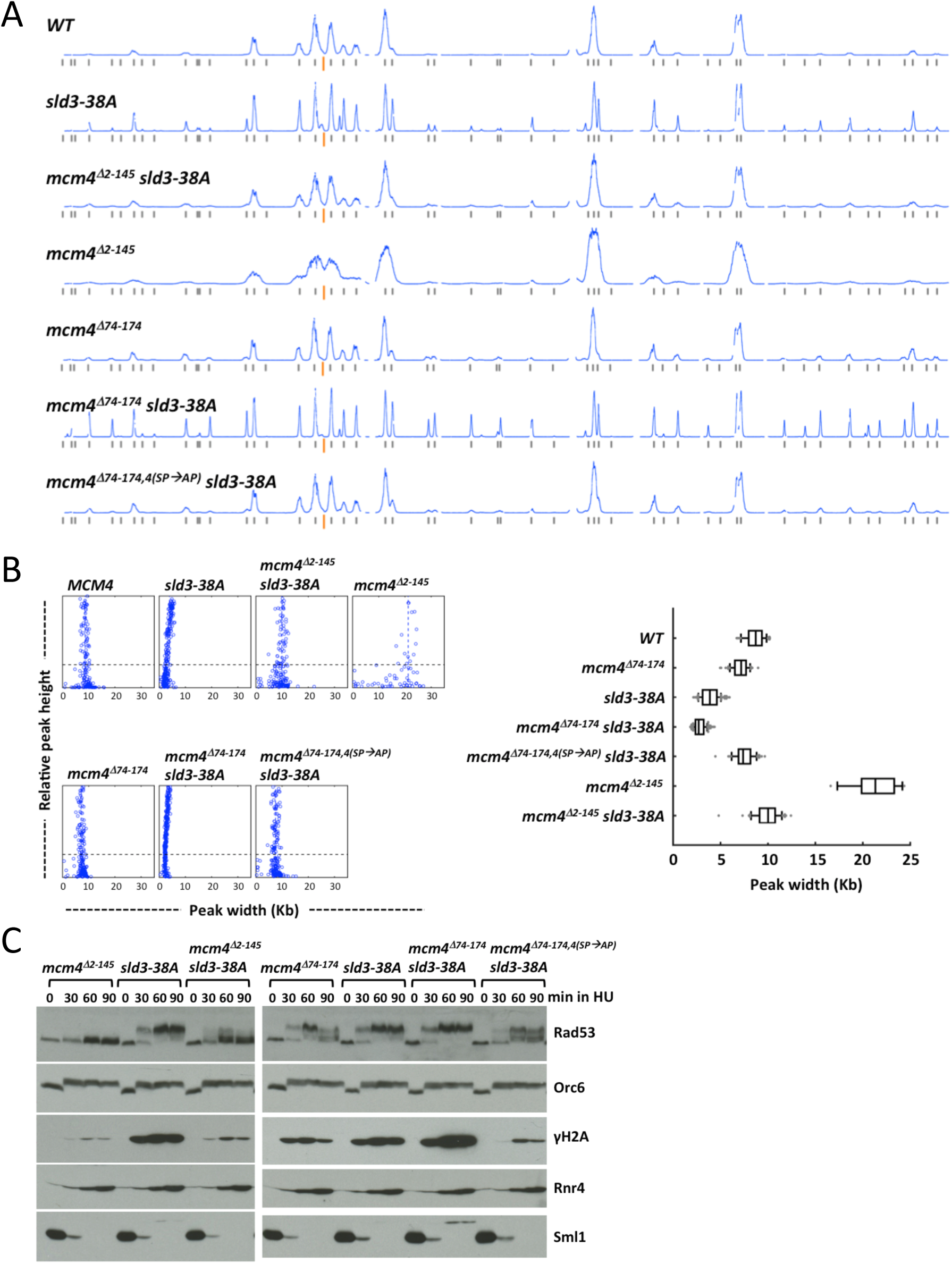
Effects of *sld3-38A* and the Mcm4 NSD distal segment on fork progression in HU. (A and B) Yeast cells were synchronized in G1 phase and released in to YPD containing 0.2 M HU and 0.5 mM EdU. (A) Replication profiles of chromosome IV for the indicated yeast strains. (B) Distribution of fork progression from origins shown as individual width-height plots and box graph, excluding peaks with heights smaller than 30 % of the maximal height scale. (C) Cells from the indicated strains were synchronized in G1, released into 0.2 M HU and collected at the indicated time points. Protein samples were analyzed as in Fig. 2C.

Hyper-activation of checkpoint signaling as measured by the degree of Rad53 and H2A phosphorylation was observed in the *sld3-38A* single mutant and the *mcm4*^*Δ74-174*^ *sld3-38A* double mutant, but was not observed in the *mcm4*^*Δ2-145*^ *sld3-38A* and *mcm4*^*Δ74-174, 4(SP→AP)*^ *sld3-38A* double mutants. However, the phosphorylation levels of the later two were slightly more elevated than that of the *mcm4*^*Δ2-145*^ single mutant (Fig. 4C). Despite the variation in Rad53 and H2A phosphorylation, the downstream response to checkpoint activation, as manifest in degradation of Sml1 and up regulation of Rnr4, appeared similar among all strains.

### Late origin firing in HU in the absence of checkpoint signaling kinases

To gain insight into the extent of late origin firing in the cells lacking the checkpoint kinases Mec1 and Rad53, the main regulators of late origin firing under genotoxic stress, we examined the replication profiles in HU in the absence of these kinases. Mec1 and Rad53 are essential for cell growth, however their essential role in growth can be bypassed by deleting the *SML1* gene, a negative regulator of the dNTP pool (Zhao et al. 1998). Thus, we analyzed *sml1Δ*, *mec1*Δ sml1Δ and *rad53*Δ *sml1*Δ cells synchronously replicating in the presence of HU for 90 min, along with the *WT* and the *mcm4*^*Δ74-174*^ *sld3-38A dbf4-19A* mutant (Fig. 5). In the *sml1*Δ cells, a significant amount of late origins were activated in HU while they were inactive in *WT* (Fig. 5A, profiles *WT* and *sml1*Δ), consistent with a previous finding (Poli et al. 2012). In *mec1*Δ *sml1*Δ and *rad53*Δ *sml1*Δ cells, more late origins were activated in HU than in the *sml1*Δ cells (Fig. 5A). However, the level of late origin firing relative to the early origins in the absence of Mec1 or Rad53 did not appear to surpass that in the *mcm4*^*Δ74-174*^ *sld3-38A dbf4-19A* triple mutant cells.

**Figure 5.**
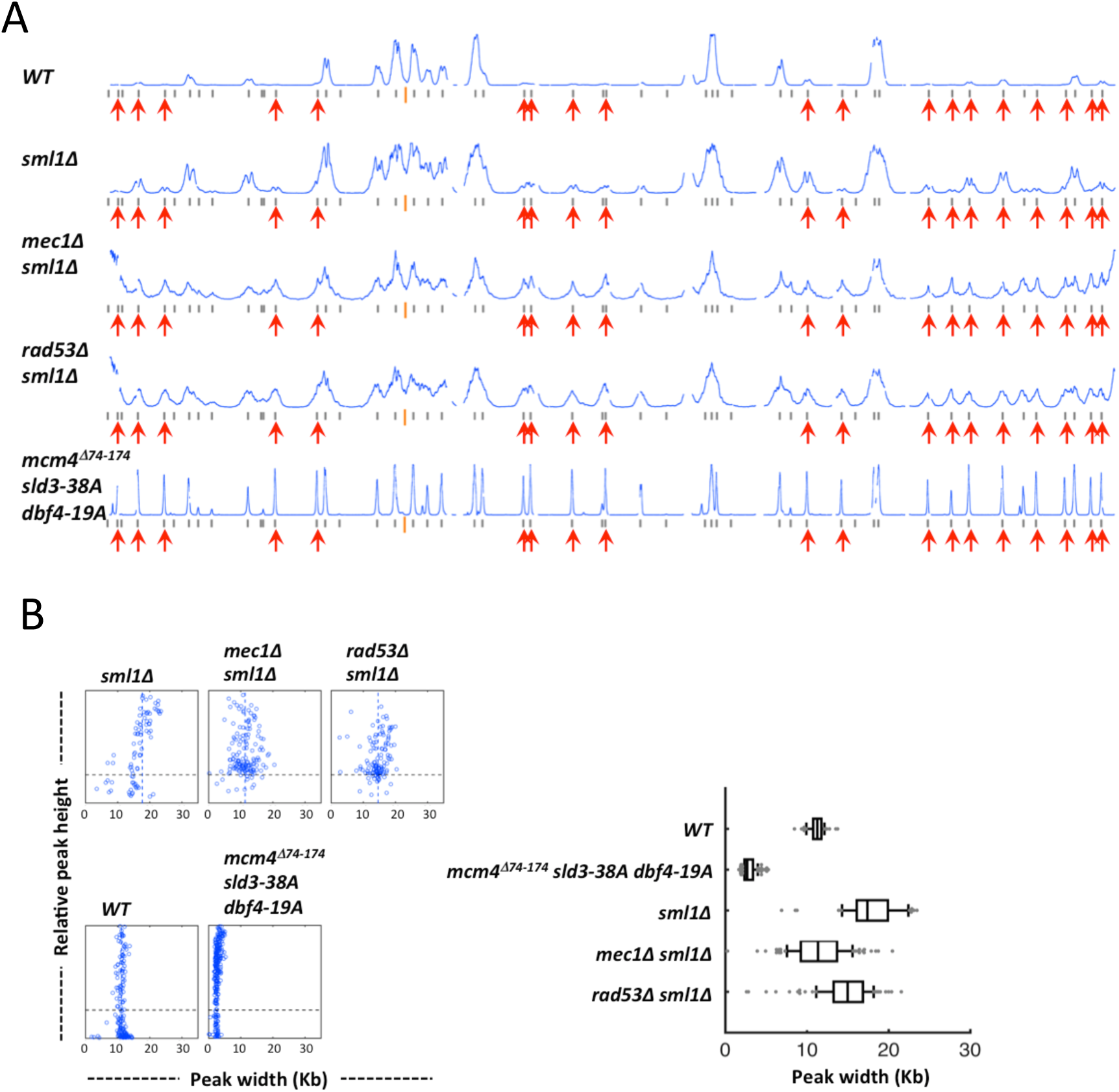
Late origin firing in HU in the absence of checkpoint signaling kinase. Yeast cells were synchronized in G1 phase and released in to YPD containing 0.2 M HU and 0.5 mM EdU for 90 min. (A) Replication profiles of chromosome IV for *WT*, *sml1Δ*, *mec1Δ sml1Δ*, *rad53Δ sml1Δ* and the *mcm4*^*Δ74-174*^ *sld3-38A dbf4-19A* triple mutant cells. (B) Distribution of fork progression from origins shown as individual width-height plots and box graph, excluding peaks with heights smaller than 30 % of the maximal height scale.

Replication forks progressed further in *sml1*Δ cells than in the *WT* in HU (Fig. 5B; (Poli et al. 2012)) because dNTP levels increased in the absence of Sml1. In *mec1Δ sml1Δ* and *rad53Δ sml1Δ* cells, replication forks progressed less than in *sml1Δ* cells, although the effect of *RAD53* deletion was moderate (Fig. 5B; Supplementary table S1). Nevertheless, it suggests that these kinases contribute to modulating fork progression. Alternatively, the lesser fork progression may reflect more aborted replication forks in the population in the absence of these kinases because they are essential for preventing DNA replication fork catastrophe under replication stress (Tercero and Diffley 2001). Importantly, fork progression in the absence of the checkpoint kinases was much greater than fork progression in the *mcm4*^*Δ74-174*^ *sld3-38A dbf4-19A* triple mutant cells.

High levels of DNA synthesis near the chromosome ends were detected in the both *mec1Δ sml1Δ* and *rad53Δ sml1Δ* cells replicating in HU, but not in *sml1Δ* cells (Fig.5A). This phenomenon is prevalent at most of the chromosome ends in these mutants. Such high level of DNA synthesis near telomeres does not occur in the unperturbed S phase (data not shown). It was not clear whether these DNA syntheses initiate from newly activated origins near telomeres. Replication profile analysis was also done with the 30 kb at each end of each chromosome masked (Supplementary figure S1 and table 1). This had an effect on the scaling of the profiles for the *mec1* and *rad53* mutants but barely affected the identification of peaks and calculation of fork progression.

### The proximal segment of the Mcm4 NSD controls firing of late origins in an unperturbed S phase

Since the proximal segment of the NSD imposes a barrier to refrain late origin firing in HU (Fig. 3B), it is possible that this domain also controls late origin firing in a normal, unperturbed S phase. To test this idea, the replication profiles of *WT* and various *mcm4* mutants containing mutations within the NSD were examined and compared at 25 min after release from G1 arrest (Fig. 6). This time point was selected because peaks representing DNA synthesis from individual origins could be clearly detected without excessive overlap and thus were more suitable for analysis of origin firing and fork progression. At this time, DNA synthesis from early origins was readily detected in *WT*, whereas very small amounts of DNA synthesis occurred from late origins (Fig. 6A, profile *WT*, red arrows). Thus, this time represents a point in the early S phase. A similar profile was found with the mutant lacking the distal segment of the NSD (Fig. 6A, profile *mcm4*^*Δ2-145*^). In *mcm4*^*Δ2-174*^ and *mcm4*^*Δ74-174*^ mutants that lacked the proximal segment of the NSD, however, peaks corresponding to firing from late origins were clearly observed, although still less than early origins (Fig. 6A), suggesting that the proximal NSD segment contributes to the temporal pattern of late origin firing during an unperturbed S phase.

**Figure 6.**
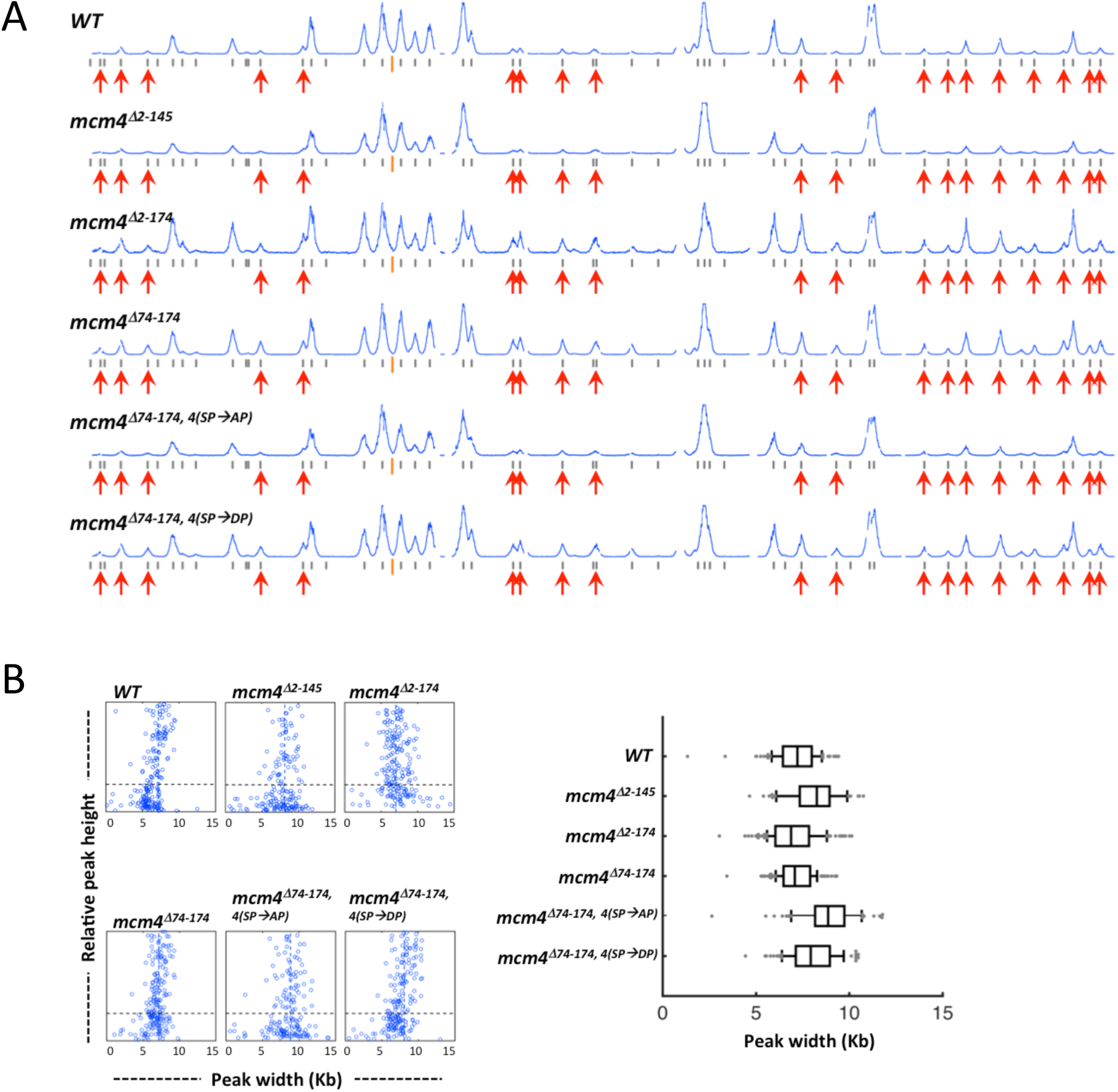
Removing the Mcm4 proximal NSD allows advanced firing of late origins in an unperturbed S phase. Yeast cells were synchronized in G1 phase and released in to YPD containing 0.5 mM EdU for 25 min at 30 ^°^C. (A) Replication profiles of chromosome IV for *WT* and Mcm4 NSD mutants. Red arrows indicate late origins that are inactive in the wild type cells but fire in the triple mutant in HU as in Fig. 2. (B) Distribution of fork progression from origins shown as individual width-height plots and a box graph, which excludes peaks with heights smaller than 30 % of the maximal height scale.

Advanced firing of late origins did not occur when, in the context of an Mcm4 lacking the proximal segment of the NSD, the phospho-acceptors for S phase-CDK phosphorylation within the distal segment of the NSD were mutated to alanine (profile *mcm4*^*Δ74-174, 4(SP→AP)*^) (Fig. 6). In contrast, mutation of these same residues to the phospho-mimetic aspartic acid (profile *mcm4*^*Δ74-174, 4(SP→DP)*^) allowed earlier firing of late origins, similar to the *mcm4*^*Δ74-174*^ mutant. Thus, phosphorylation of the CDK sites within the distal segment of the NSD is important for efficient firing of late origins during an unperturbed S phase.

The replication profiles of *sld3-38A* and *dbf4-19A* mutants were examined in an unperturbed S phase (Fig. 7). None of the *sld3-38A*, *dbf4-19A* and *sld3-38A dbf4-19A* mutants showed advanced firing of late origins compared to the replication profile of wild type cells (Fig. 7A). Furthermore, combining *mcm4*^*Δ74-174*^ with either *sld3-38A* or *dbf4-19A* mutations, or even the triple mutant *mcm4*^*Δ74-174*^ *sld3-38A dbf4-19A* did not lead to more firing of late origins than the *mcm4*^*Δ74-174*^ single mutant (Fig 7A). Thus, unlike the NSD proximal segment mutant, checkpoint kinase Rad53 phosphorylation of Sld3 or Dbf4 does not control late origin firing during a normal S phase.

**Figure 7.**
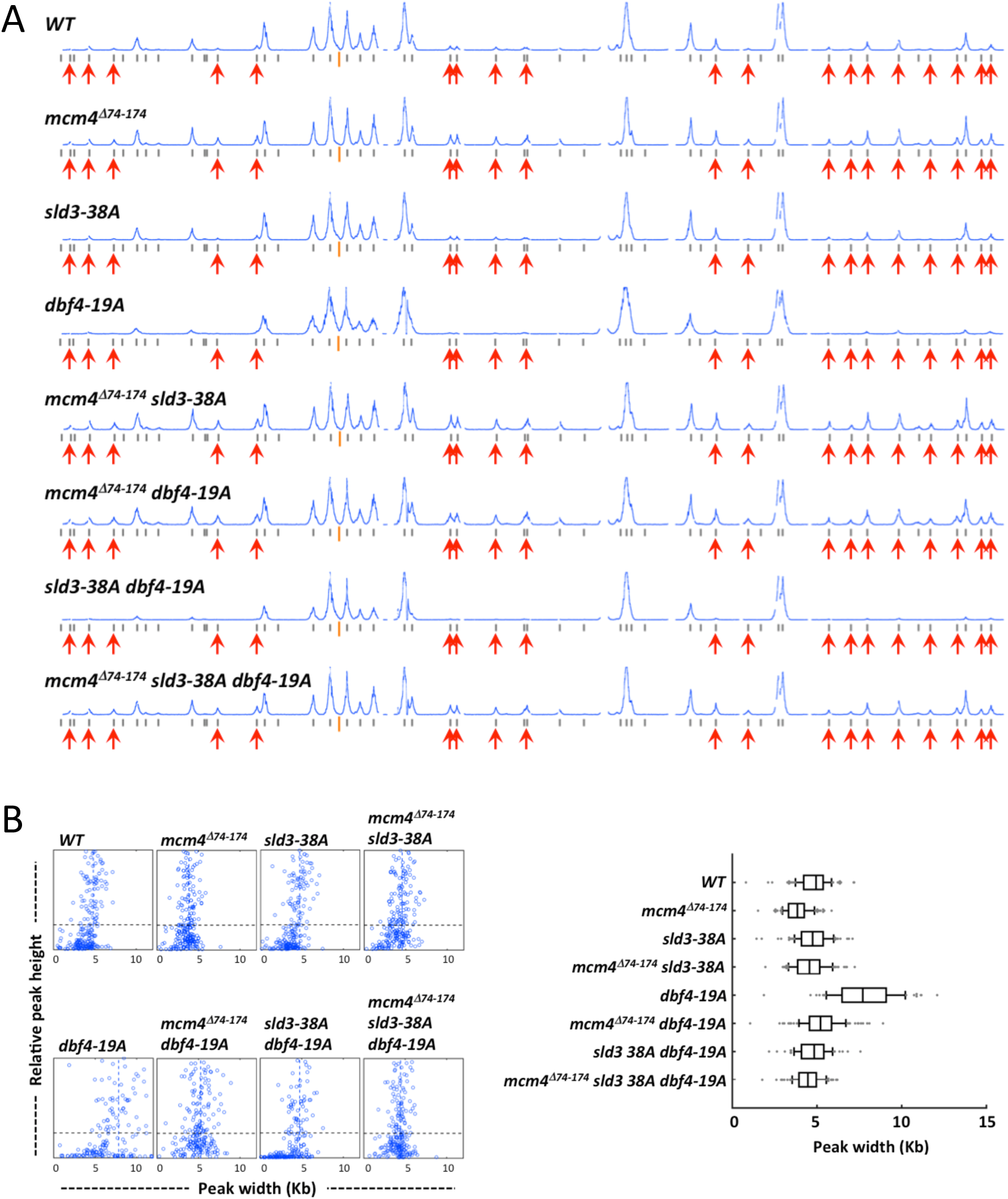
Checkpoint resistant mutations of *SLD3* and *DBF4* do not affect timing of late origin firing in an unperturbed S phase. Yeast cells were synchronized in G1 phase and released into YPD containing 0.5 mM EdU for 25 min. (A) Replication profiles of chromosome IV for *WT*, single, double and triple mutants with *mcm4*^*Δ74-174*^, *sld3-38A* and/or *dbf4-19A* alleles as indicated on the top-left of each profile. (B) Distribution of fork progression constructed as in Fig. 4B.

From the replication profile analyses, it appeared that mutations in the Mcm4 NSD did not have a dramatic effect on fork progression in an unperturbed S phase (Fig. 6B; Supplementary table S1). However, subtle differences would be more difficult to detect in such an experiment because DNA synthesis occurred much faster in the absence of HU. Similarly, the *sld3-38A* mutation did not restrict fork progression in contrast to its effect in HU (Fig. 7B; Supplementary table S1). The distribution of fork progression in the *dbf4-19A* mutant appeared more heterogeneous (Fig. 7B), similar to the pattern observed in HU (Fig. 2B). Double mutants of *dbf4-19A* with either *mcm4*^*Δ74-174*^ or *sld3-38A* yielded phenotypes resembling the single mutant of *mcm4*^*Δ74-174*^ or *sld3-38A* with respect to replication fork progression.

### Checkpoint response and replication profiles for Mcm4 NSD mutants entering S phase in the presence of MMS

Although the NSD proximal segment controls late origin firing in both a normal S phase and S phase with a depleted pool of dNTP, the effect of NSD mutations on fork progression was only observed in the presence of HU. Furthermore, *sld3-38A* showed a strong phenotype in restricting fork progression in HU, but no obvious effect in a normal S phase. The differential influence of these mutations on origin firing and replication fork progression in an unperturbed versus a HU treated S phase raised the question of whether these factors would have same effect on other types of genotoxic stress than dNTP depletion. Thus, the DNA damage checkpoint response and DNA replication profiles in cells replicating in the presence of the DNA alkylating agent methyl methanesulfonate (MMS) were studied.

For replication profile analyses, cells were synchronized in G1 phase and allowed to enter S phase in the presence of MMS for 50 min. We did not use the 90 min time point that we typically used for analysis of profile in HU because replication in MMS was faster than in HU and 90 min in MMS would have produced profiles difficult to analyze due to numerous merged peaks and passive replication at late/unfired origin loci by replication forks moving from early firing origins. At 50 min in MMS after release from G1 arrest, few late origins fire in *WT* and in the mutant lacking the distal NSD segment (Fig. 8A, profiles *WT* and *mcm4*^*Δ2-145*^). In contrast, late origin firing was more evident in the mutants lacking the proximal NSD segment (Fig. 8A, profiles *mcm4*^*Δ2-174*^ and *mcm4*^*Δ74-174*^). In the same context, mutation of the CDK sites to alanines within the distal NSD segment (*mcm4*^*Δ74-174,4(SP→AP)*^) suppressed late origin firing while mutating the same sites to phosphomimetic aspartic acids (*mcm4*^*Δ74-174,4(SP→DP)*^) restored the level of late origin firing. Thus, the proximal NSD segment also mediates control of late origin firing in MMS.

**Figure 8.**
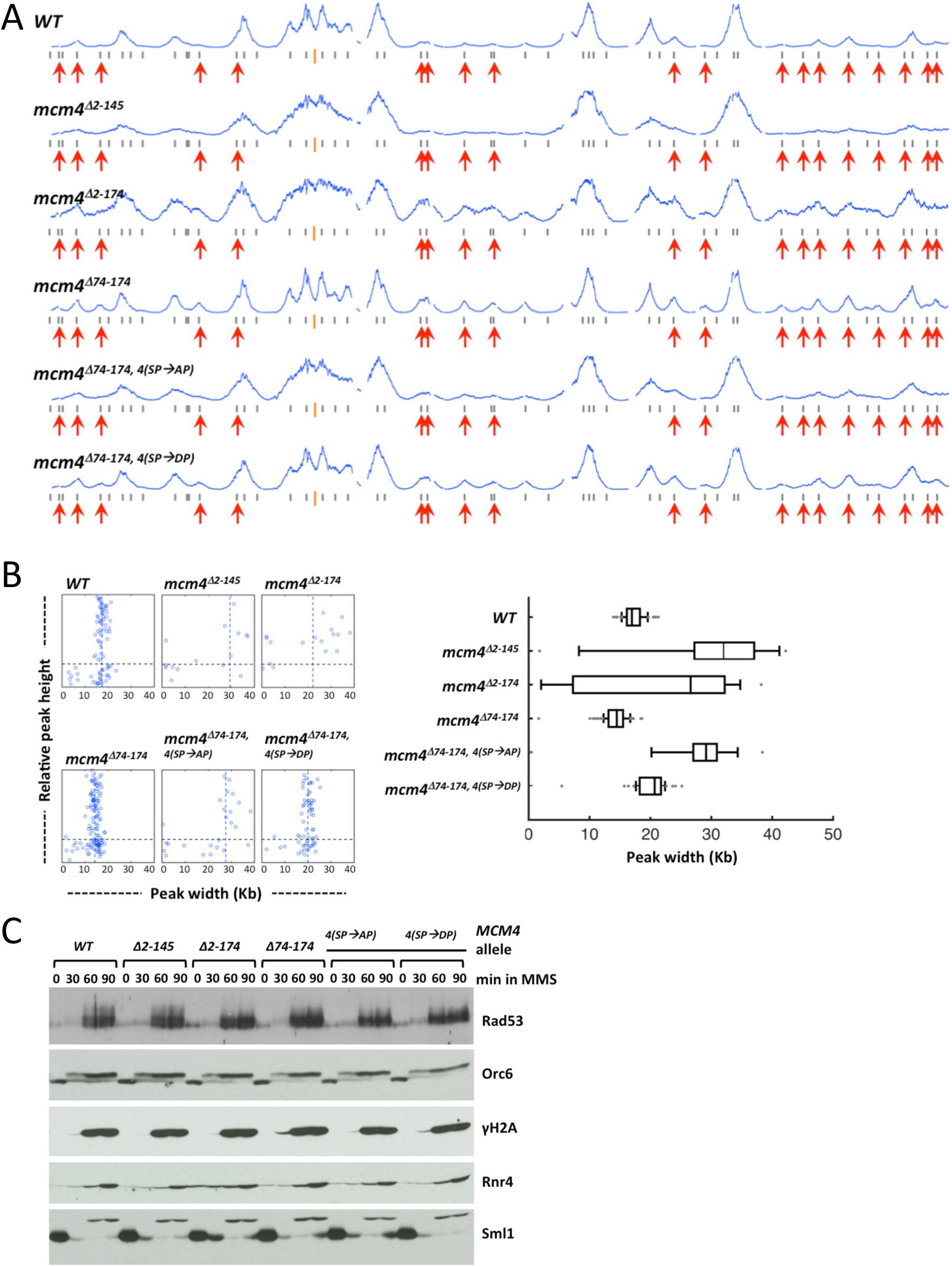
Replication profiles and checkpoint response in Mcm4 NSD mutants. (A and B) Yeast cells were synchronized in G1 phase and released into YPD containing 0.05 % MMS and 0.5 mM EdU for 50 min. (A) Replication profiles of chromosome IV for the wild type and the indicated Mcm4 NSD mutants. (B) Distribution of fork progression from origins shown as individual width-height plots and a box graph, which excludes peaks with heights smaller than 30 % of the maximal height scale. (C) Cells from the indicated strains were synchronized in G1, released into YPD containing 0.05 % MMS and collected at indicated time points. Protein samples were prepared using TCA extraction and analyzed by immunoblot.

Replication fork progression was also affected in the Mcm4 NSD mutants replicating in the presence of MMS (Fig. 8). Fork progression was more restricted in the *mcm4*^*Δ74-174*^ mutant lacking the NSD proximal segment compared to *WT*, while more expansive fork progression was observed in mutants lacking the distal NSD segment (Fig. 8A and B; compare *mcm4*^*Δ2-145*^ with *WT* and *mcm4*^*Δ2-174*^ with *mcm4*^*Δ74-174*^). Fork progression in MMS was also regulated by phosphorylation at the CDK target sites within the distal segment of the NSD because forks progressed further in *mcm4*^*Δ74-174,4(SP→AP)*^ compared to progression in *mcm4*^*Δ74-*174^, while in the *mcm4*^*Δ74-174,4(SP→DP)*^ mutant fork progression was more restricted, similar to that in *mcm4*^*Δ74-*174^ (Fig. 8B). Therefore, the distal and proximal segments of the Mcm4 NSD play important roles in mediating control of fork progression in diverse types of genotoxic agents.

In HU, the NSD distal segment was important for checkpoint signaling at the level of Mec1 signaling (Sheu et al. 2014), although Mec1 phosphorylation of Mcm4 is independent of checkpoint activation (Randell et al. 2010). Removing the distal segment of the NSD, or mutation of the phospho-acceptor amino acids at CDK sites to alanine within this domain resulted in reduced levels of Rad53 hyper-phosphorylation and γH2A. In contrast, other aspects of checkpoint signaling further downstream, such as Sml1 degradation and Rnr4 induction appeared normal in cells treated with HU. In MMS, however, hyper-phosphorylation of Rad53 and S129 phosphorylation in H2A, as well as further downstream events such as degradation of Sml1 and up regulation of Rnr4 levels appeared very similar among wild type and various Mcm4 NSD mutants (Fig. 8C). Thus, the Mcm4 NSD did not play a prominent role in checkpoint signaling in response to DNA damage caused by MMS.

### Cooperation between the proximal segment of the Mcm4 NSD and Rad53 in regulating late origin firing in MMS

The Rad53 resistant *sld3-38A* and *dbf4-19A* strains alone also showed low levels of late origin firing in cells treated with MMS (Fig. 9A). The difference between the wild type and the *dbf4-19A* mutant was subtle. Nevertheless, the double mutants *sld3-38A dbf4-19A*, *mcm4*^*Δ74-174*^ *sld3-38A* and *mcm4*^*Δ74-174*^ *sld3-38A* activated more late origins than the wild type and any of the single mutants. The *mcm4*^*Δ74-174*^ *dbf4-19A* showed only a slight increase in late origin firing, compared with the *mcm4*^*Δ74-174*^ single mutant, consistent with Mcm4 functioning downstream of Dbf4 in controlling late origin firing in MMS. Furthermore, the triple mutant activated late origins the most efficiently (Fig. 9A, profile *mcm4*^*Δ74-174*^ *sld3-38A dbf4-19A*). Thus, all three factors contribute to control of late origin firing through overlapping but non-identical pathways in MMS, as was found for cells treated with HU.

**Figure 9.**
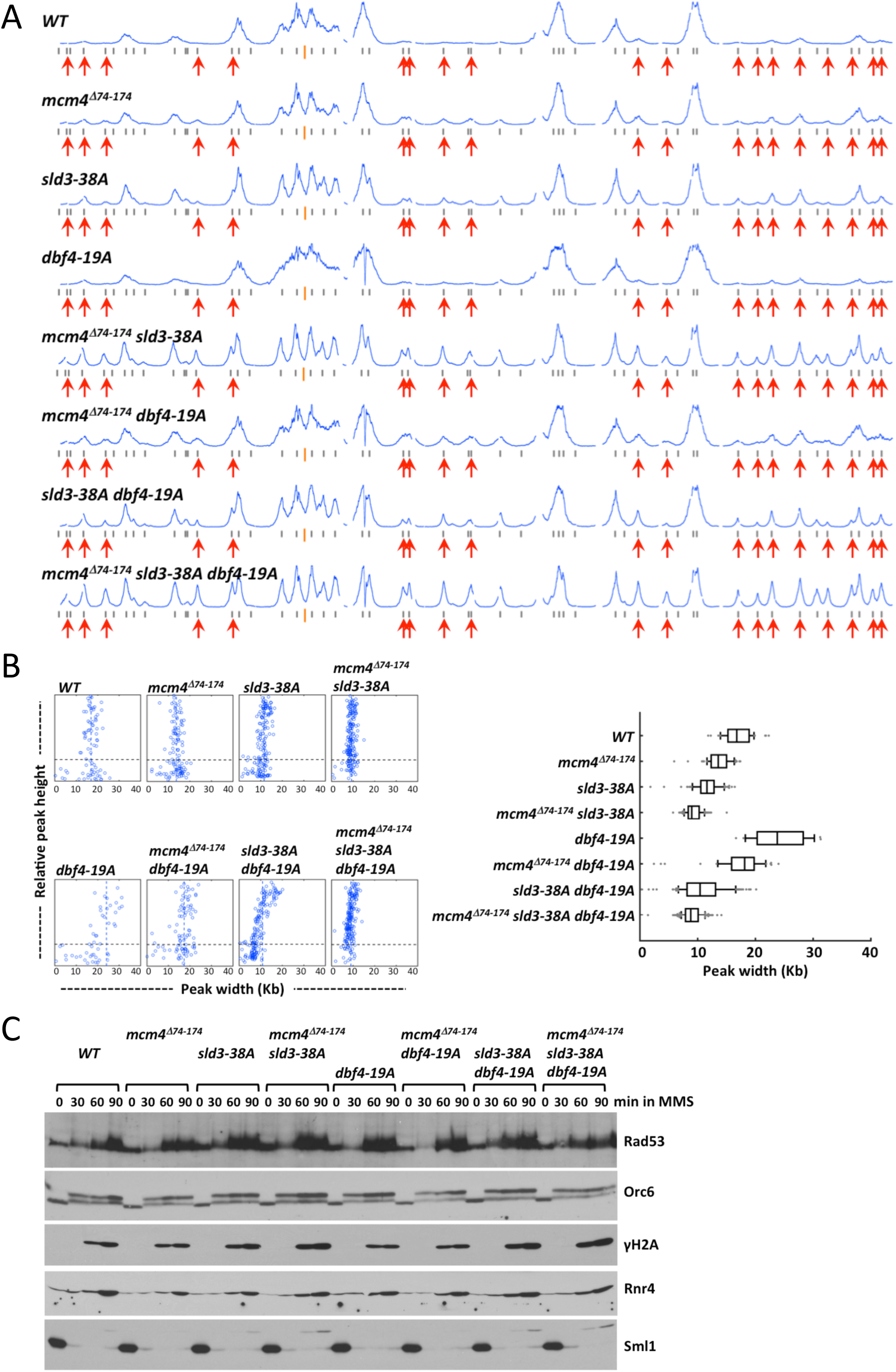
Analysis of replication profiles and checkpoint response in the wild type, single, double and triple mutants with *mcm4*^*Δ74-174*^, *sld3-38A* and/or *dbf4-19A* alleles. (A and B) Yeast cells were synchronized in G1 phase and release into YPD containing 0.05 % MMS and 0.5 mM EdU for 50 min. (A) Replication profiles of chromosome IV. (B) Distribution of fork progression constructed as in Fig. 4B. (C) Cells from the indicated strains were synchronized in G1, released into YPD containing 0.05 % MMS and analyzed as in Fig. 8C.

Like in HU, the *sld3-38A* mutant had an effect on restricting replication fork progression in MMS (Fig. 9B). The effect of *sld3-38A* and *mcm4*^*Δ74-174,*^on restricting fork progression was additive under this condition, suggesting that they control fork progression separately. Fork progression in *dbf4-19A* was the more extensive, but become more restricted when *sld3-38A* and *mcm4*^*Δ74-174*^ were also present. Thus, Mcm4, Sld3 and Dbf4 cooperate to regulate fork progression in MMS.

The DNA damage checkpoint signaling was active in the wild type and all of the single, double and triple mutant combinations of *mcm4*^*Δ74-174*^, *sld3-38A* and *dbf4-19A* (Fig. 9C). Although we detected elevated H2A S129 phosphorylation in the double mutants of *mcm4*^*Δ74-174*^ *sld3-38A* and *sld3-38A dbf4-19A*, as well as the triple mutant, the differences in the signaling at this level among strains did not appear as dramatic as those observed in HU (Fig. 2C).

### Suppression of the *temperature-sensitive* (*ts*) phenotype of multiple *sld3-ts* mutants by deletion of the Mcm4 NSD proximal segment

Although replication profile analyses suggest that Sld3 and the Mcm4 NSD mediate controls from separate pathways, the fact that they affect similar processes raises the possibility that the tasks executed by these two factors may converge on a common target. For example, the proximal segment of the Mcm4 NSD maybe inhibiting the same molecular process that Sld3 is facilitating. If this is the case, it is likely that removing the proximal NSD segment would compensate for the weakened Sld3 function in hypomorph *sld3* mutants. The idea was tested by introducing the *mcm4*^*Δ74-174*^ mutation in *sld3-ts* mutants *sld3-5*, *sld3-6* and *sld3-7* (Kamimura et al. 2001), all of which fail to grow on YPD plates at a non-permissive temperature above 30°C, 37°C and 34°C, respectively. At 30°C, the *sld3-5* mutant grew extremely poorly, compared with the wild type and the *mcm4*^*Δ74-174*^ mutant, while the *mcm4*^*Δ74-174*^ *sld3-5* double mutant grew much better than the *sld3-5* mutant (Fig. 10A). At 23°C, the *sld3-5* mutant also grew slower than the wild type but the *mcm4*^*Δ74-174*^ *sld3-5* grew similarly to the wild type. Thus, removing proximal segment of the NSD improved the growth of the *sld3-5* mutant. Likewise, removing the proximal segment of the NSD improved the growth of the *sld3-6* and *sld3-7* mutants at 37°C, but no growth occurred in *sld3-5* and *mcm4*^*Δ74-174*^ *sld3-5* at this temperature. Thus, removing the Mcm4 proximal segment of the NSD suppresses the defect of multiple hypomorph *sld3-ts* mutants.

**Figure 10.**
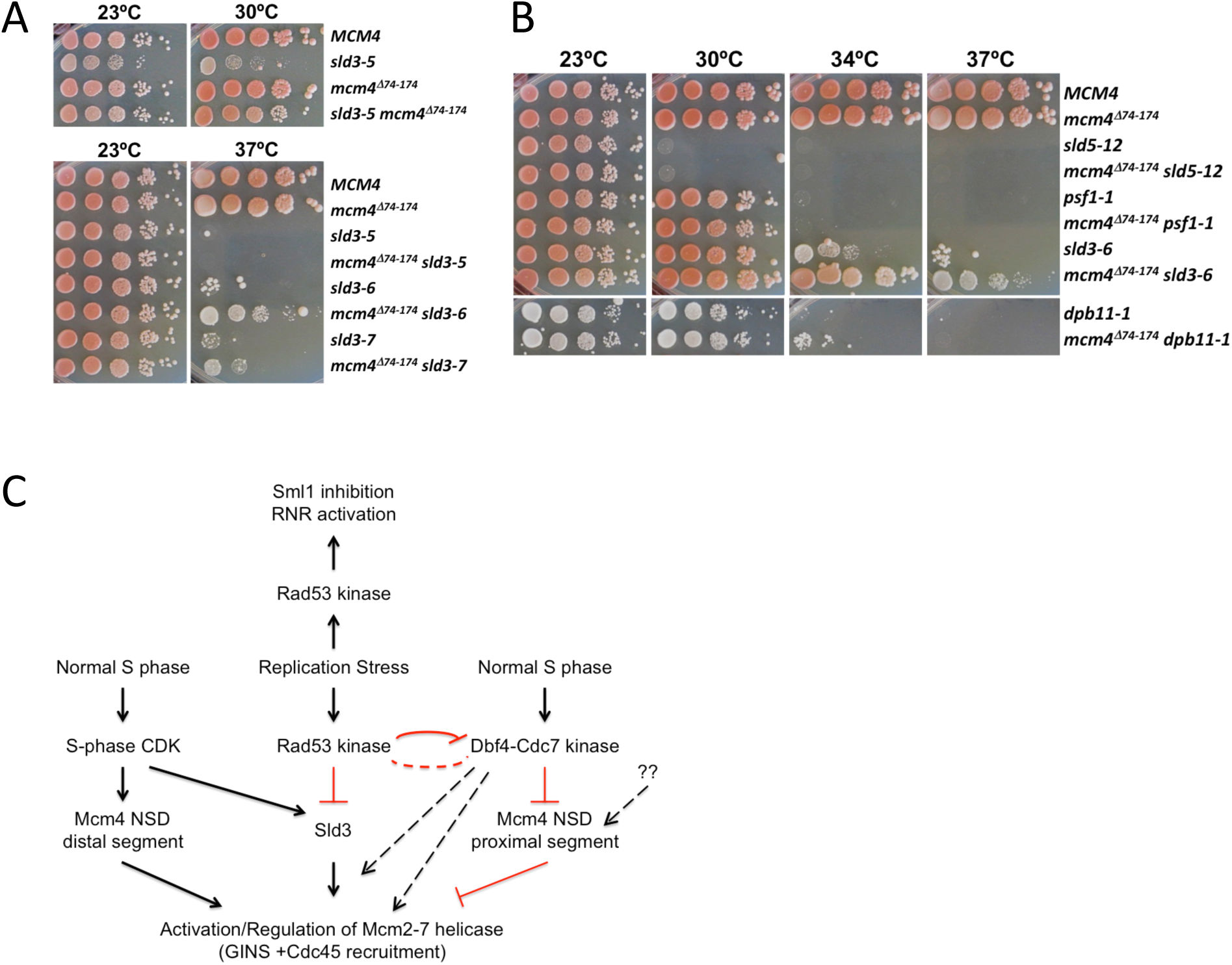
Partial suppression of the sld3-ts by the *mcm4*^*Δ74-174*^ mutant. (A and B) Serial 10-fold dilution of 10^5^ yeast cells on YPD at the indicated temperatures. (C) A model for control of Mcm2-7 helicase.

We also tested if removing the proximal segment of the Mcm4 NSD would also suppress the *ts* phenotype of mutants affecting other factors that function with Sld3, such as Dbp11 and components of the GINS complex (Kamimura et al. 1998; Takayama et al. 2003). The *mcm4*^*Δ74-174*^ mutation failed to suppress the *ts* phenotype of *sld5-12* or *psf1-1*, mutants in GINS subunits (Fig. 10B, 30°C, 34°C and 37°C). A very slight improvement of growth in the *mcm4*^*Δ74-174*^ *dpb11-1*, compared with the *dpb11-1* mutant, at 34°C was observed. However, the suppression was much less effective than the suppression of the *sld3-6* defect by *mcm4*^*Δ74-174*^ (Fig. 10B, 34°C and 37°C). The specific and strong suppression of *sld3-ts* by *mcm4*^*Δ74-174*^ is consistent with the idea that Sld3 and the Mcm4 NSD regulate the same process at the molecular level to control origin firing and influence replication fork progression under genotoxic stress.

## DISCUSSION

The inhibition of DNA replication under genotoxic stress requires both Rad53 and Mec1 kinases (Sanchez et al. 1996; Santocanale and Diffley 1998; Zegerman and Diffley 2009). In previous studies, we demonstrated that even in the presence of an active S-phase checkpoint response, late origins fire in the presence of HU when the Mcm4 NSD proximal segment was removed (Sheu et al. 2014). The observation suggested that under replication stress the checkpoint kinase Rad53 inhibited Dbf4 by phosphorylation (Lopez-Mosqueda et al. 2010; Zegerman and Diffley 2010), rendering DDK incapable of relieving the initiation inhibitory activity of the Mcm4 NSD proximal segment (Fig. 10C). Since the Mcm4 NSD proximal segment is targeted by DDK (Sheu and Stillman 2010), inhibition of DDK by active Rad53 would not prevent initiation in the absence of this initiation inhibitory domain. However, late origin firing in the absence of this domain was still rather inefficient, presumably because checkpoint activation of Rad53 still allowed phosphorylation of the other target, Sld3, thereby inactivating Sld3 activity and preventing robust firing of late origins (Fig. 10C). By using specific probes for analysis by alkaline gel electrophoresis or two-dimensional gel electrophoresis, some firing of certain late origins was detected in *sld3* and *dbf4* mutants that are refractory to the inhibition by the checkpoint kinase Rad53 (Lopez-Mosqueda et al. 2010; Zegerman and Diffley 2010). In the current study, whole genome replication profile analysis was used to investigate the individual roles, as well as the combined effect of the two Mcm4 NSD segments and the Rad53 targets, Sld3 and Dbf4, on origin activation and replication fork progression in order to delineate the relationship among these factors in control of replication in response to replication stress.

In both HU and MMS, late origins fire in each of the *mcm4*^*Δ74-174*^, *sld3-38A* and *dbf4-19A* single mutants across the entire genome, albeit very inefficiently (Fig. 2 and 8). The fact that all the double mutant combination among these three mutations activated late origins more efficiently than the respective single mutants and that the triple mutant exhibited the most efficient firing of late origins suggests that each factor contributes a unique function in control of origin activation. Yet, their functions may not be completely independent (Fig. 10C). For example, late origin firing in the *mcm4*^*Δ74-174*^ *dbf4-19A* double mutant appeared only marginally more efficient than the *mcm4*^*Δ74-174*^ single mutant. This was not surprising given that Mcm4 is the essential target of DDK and *mcm4*^*Δ74-174*^ can bypass the regulation of the kinase (Sheu and Stillman 2010). Nevertheless, because the triple mutant promotes robust firing of late origins, more than any of the single and double mutants, both the Mcm4 NSD proximal segment and Dbf4 must also independently contribute to regulate late origin firing (Fig. 10C). This can be anticipated for Dbf4 because DDK also phosphorylates other factors in addition to Mcm4 NSD. For Mcm4, it raises the possibility that, in addition to DDK, other factors might participate in the regulation of the function of the proximal NSD in controlling late origin firing under replication stress. Identification of factors that interact with the proximal NSD may shed light on this aspect of the control mechanism. Alternatively, Dbf4 may affect Mcm2-7 helicase activity independent of the Mcm4 NSD or participate in feedback regulation of Rad53 kinase activity or specificity (Fig. 10C, dashed lines). One possibility is that they antagonize each other’s activity, essentially creating a feedback loop for inactivating the checkpoint once the replication stress has subsided.

The partial overlap of functional pathways involving Sld3 and Dbf4, as revealed by the minimal combined effect of *dbf4-19A* and *sld3-38A* on late origin firing in HU compared to each single mutant can be explained by the fact that the early association of Sld3 to the pre-RC depends on DDK activity (Heller et al. 2011; Tanaka et al. 2011) (Fig. 10C). In contrast, the *mcm4*^*Δ74-174*^ *sld3-38A* exhibited the strongest additive effect among the double mutant combinations, suggesting that these two factors mediate regulation via separate pathways (Fig. 10C). However, the suppression of the hypomorph *sld3-ts* defect by *mcm4*^*Δ74-174*^ suggests that these two pathways intersect on a common process and they are likely regulating the same factors. It is possible that this common pathway merges on the direct activation of the Mcm2-7 helicase by recruiting other helicase components Cdc45 and GINS (Fig. 10C).

In the *sld3-38A dbf4-19A* double mutant that was expected to be completely refractory to the control by the Rad53-dependent S-phase checkpoint, late origins did not fire until 50 min into S phase in the presence of HU (Fig. 3B). Given that late origin firing was readily detected in a normal, unperturbed S phase at this time, this suggests that a mechanism is functioning to withhold late origins from firing in this double mutant condition. Removing the proximal segment of the Mcm4 NSD in the same genetic background allowed late origins to fire by 25 min after release, similar to what we have observed in unperturbed S phase (Fig. 6A), strongly suggesting that activating DDK alone is not sufficient to efficiently block the function of the proximal NSD segment in withholding late origin from firing earlier in HU. Thus, besides DDK, additional factors might participate in relieving the block imposed by proximal NSD to delay late origin firing.

The proximal segment of the Mcm4 NSD also controls late origin firing in an unperturbed S phase (Fig. 10C). In mutants lacking this domain, more late origin firing was detected in early S phase (Fig. 6A). In contrast, neither of the *sld3-38A*, *dbf4-19A* or *sld3-38A dbf4-19A* mutants showed advanced firing of late origins in an unperturbed S phase (Fig. 7A). Thus, checkpoint kinase Rad53 does not appear to control late origin firing through Sld3 and Dbf4 in a normal S phase. Phosphorylation of the CDK sites within the distal segment of the Mcm4 NSD (Devault et al. 2008) was also important for advanced firing of late origins in the unperturbed S phase when the proximal segment of the NSD was removed (Fig. 6A). Previous studies in budding yeast have shown that, in the absence of the main S-CDK cyclin, Clb5, only early origins fire, but not late origins (Donaldson et al. 1998). Thus, activation of late origins requires activity of S phase CDK. Together, these results suggest that phosphorylation the distal segment of the Mcm4 NSD by CDK is an important step for activation of late origins (Fig. 10C). The accumulation of CDK activity as cells progress through S phase may eventually allow late origins to fire.

The role of the Mcm4 NSD in regulating late origin firing and fork progression previously discovered in HU was largely recapitulated in cells replicating in MMS (Fig. 8). Specifically, in MMS the proximal segment of the NSD mediates control of late origin firing and the distal segment NSD mediates control of fork progression in a manner that is regulated by phosphorylation at CDK target sites. However, unlike the response in HU, the Mcm4 NSD mutations exhibited little effect on checkpoint signaling in response to DNA damage caused by MMS (Fig. 8C). In our previous study, we noticed an inverse correlation between checkpoint signaling and DNA replication fork progression in HU (Sheu et al. 2014), raising the possibility that one process controls the other. The study here in MMS, in contrast, provides evidence that these two processes might not always influence each other. At least in MMS, the Mcm4 NSD is likely to regulate fork progression through a mechanism independent of the canonical DNA damage checkpoint pathway. One possibility is that the DNA damaging signal would somehow control the activity of CDK or other SP site kinases, which in turn regulate the function of the distal segment of the NSD.

Sld3 also mediates control of DNA replication fork progression in both HU and MMS (Fig. 2B and 8B). The *sld3-38A* mutant exhibits a dramatic slowdown in replication fork progression. This is a somewhat surprising observation because Sld3 is not considered a replication fork component because it is required for initiation, but not for elongation in normal S phase progression (Kanemaki and Labib 2006). It is not clear why DNA synthesis is so limited in the this mutant but the mutation does not seem to result in a defective replication factor because the replication profile of this mutant is similar to the wild type in an unperturbed S phase and this mutant grows at a rate comparable to wild type cells. Furthermore, a previous study reported that yeast strains expressing *sld3-38A* as the sole copy of Sld3 displayed no increase in sensitivity to hydroxyurea or DNA damaging agents and did not exhibit synthetic growth defects with several conditional alleles of essential replication proteins (Zegerman and Diffley 2010). Therefore, this phenotype is likely due to regulation of fork function. Replication profile analysis of the double mutant combining *sld3-38A* and NSD mutants in the distal segment showed that the effect of the *sld3-38A* mutant on fork progression in HU and distal NSD mutant are not epistatic to each other (Fig. 4), consistent with the idea that these two factors operate in separate pathways to regulate fork progression in HU. In contrast to its role in HU and MMS, the checkpoint resistant *sld3-38A* mutant did not affect fork progression in an unperturbed S phase (Fig. 7). Thus, the control of fork progression through Rad53 target sites on Sld3 is a specific feature in the genotoxic-stressed condition.

It has been reported that the level of origin firing inversely affected the rate of fork progression (Zhong et al. 2013) presumably because active forks compete for limiting dNTP pools, which is an important determinant of fork progression (Poli et al. 2012). Since Mcm4 NSD, Sld3 and Dbf4 mutation increased the number of origins that were activated, it was possible that their effects on replication fork progression in HU were in part the consequence of competition for limiting dNTP pools or limiting DNA replication proteins. We suggest, however, that the limited replication fork progression in the *sld3-38A* and *mcm4*^*Δ74-174*^ mutants was a result of these proteins directly involved in controlling replication fork progression and not only due to more active origins competing for limiting dNTP or replication proteins since in the *rad53* and *mec1* mutants, late origins were equally active but replication fork progression was much greater than fork progression in the *sld3-38A* and *mcm4*^*Δ74-174*^ mutants. Furthermore, MMS treatment should not affect dNTP levels like addition of HU, but the effects of *sld3-38A* and *mcm4*^*Δ74-174*^ mutations on replication fork progression in MMS paralleled the effects observed in HU. Thus, although increased origin firing may correlate with slowing of DNA replication fork progression, the level of origin firing may not be the sole explanation for changes we observe in replication fork progression.

Since DDK binds directly to the MCM-2-7 helicase subunits Mcm4 and Mcm2 (Varrin et al. 2005; Sheu and Stillman 2006; Jones et al. 2010), and DDK binds directly to Rad53 (Dohrmann et al. 1999; Weinreich and Stillman 1999; Kihara et al. 2000), it is possible that the regulation of the response to DNA replication stress such as limiting dNTP levels involves a local response at the DNA replication fork, essentially a solid state regulatory complex. How other Mcm2-7 associated replication checkpoint proteins such as Mrc1, Dpb11, Sld2 and the large subunit of DNA polymerase ε control initiation of replication and fork progression remains to be investigated, but we suspect that they integrate with the regulatory system involving Dbf4, Sld3, and Mcm4.

## METHODS

### Yeast strains and methods

Yeast strains generated in this study were derived from W303-1a (*MAT****a*** *ade2-1 can1-100 his3-11,15 leu2-3,112 trp1-1 ura3-1*) and are described in Supplementary table 2. A two-step gene replacement method was used to replace the endogenous *MCM4* with *mcm4* mutants as described (Sheu et al. 2014). All the yeast strains used for the whole genome DNA replication profile analyses have a copy of the BrdU-Inc cassette inserted into the *URA3* locus (Viggiani and Aparicio 2006). For G1 arrest of *bar1 Δ* strains, exponentially growing yeast cells (∼10^7^ cell/ml) in YPD were synchronized in G1 with 25 ng/ml of α-factor for 150 min at 30°C. For G1 arrest of *BAR1* strains, exponentially growing cells were grown in normal YPD, then transferred into YPD (pH3.9), grown to approximately 10^7^ cell/ml and then synchronized in G1 with three doses of α-factor at 2 μg/ml at 0, 50, and 100 min time point at 30°C. Cells were collect at 150 min for release. To release from G1 arrest, cells were collected by filtration and promptly washed twice on the filter using 1 culture volume of H_2_O and then resuspended into YPD medium containing 0.2 mg/ml pronase E (Sigma).

### Protein sample preparation and immunoblot analysis

TCA extraction of yeast proteins was as described previously (Sheu et al. 2014). For chromatin fractionation, chromatin pellets were prepared from ∼5×10^8^ yeast cells and chromatin-bound proteins were released using DNase I using a procedure described previously (Sheu et al. 2014). For immunoblot analysis, proteins samples were fractionated by SDS-PAGE and transferred to a nitrocellulose membrane. Immunoblot analysis for Mcm3, Cdc45, Orc6, Mcm4, Rad53, γ-H2A, Rnr4 and Sml1 were performed as described (Sheu et al. 2014).

### Isolation and preparation of DNA for whole genome replication profile analysis

Detailed protocol was described previously (sheu2014). Briefly, yeast cells were synchronized in G^1^ with α-factor and released into medium containing of 0.2 mg/ml pronase E, 0.5 mM 5-ethynyl-2’-deoxyuridine (EdU) with or without addition of 0.2 M HU or 0.05% MMS as described in the main text. At the indicated time point, cells were collected for preparation of genomic DNA. The genomic DNA were fragmented and then ligated to adaptors containing custom barcodes and then biotinylated, purified, PCR-amplified, quantified, pooled and submitted for sequencing. Computational analyses of sequencing data were described in detail previously (Sheu et al. 2014).

### Computational analyses of sequencing data

Read mapping, replication profile analysis, and peak width analysis were performed as previously described (Sheu et al. 2014) with minor modifications. Supplemental table 1 lists the specific analysis parameters used for each sample. Briefly, each genome-wide replication profile was generated from between 4.4 to 49.2 million reads mapped to the sixteen *Saccharomyces cerevisiae* S288C chromosomes (NC_001133 through NC_001148). As in (Sheu et al. 2014), only the first 31bp of each read were used for mapping. Read counts were then averaged across the genome using a sliding window of 500 bp, 1000 bp, or 2000 bp. Genomic positions that could not be reliably mapped to themselves in a way that substantially affected these smoothed profiles were masked. Positions 450000:500000 of chromosome XII, which encompass the rDNA locus, were also masked. In the specific analyses performed for Figure S1, 30 kb at each end of each chromosome was masked as well. The replication profiles for each data set were then rescaled so that profile height was less than or equal to one “unit” at 99.5 percent of genomic positions. This rescaling was solely to facilitate the visual comparison of profiles and had no effect on subsequent peak width quantification. Peak widths were quantified using the full width at half maximum of each replication profile peak that encompassed a single annotated origin in oriDB v2.1.0 (Siow et al. 2012). Peaks that encompassed multiple origins or masked genomic regions were discarded. Peaks having a height below either 10% or 30% of the “unit” value were also discarded in order to avoid the analysis of widths from insufficiently well-defined peaks. This height cutoff is indicated by a line in each of the peak-width plots in the Figures.

All DNA sequencing data was deposited on the NCBI sequence read archive (accession number SRA279689; http://www.ncbi.nlm.nih.gov/sra/?term=SRA279689). The data analysis scripts are publically available on Github: https://github.com/jbkinney/14_sheu

## ACKNOWLEDGMENTS

This work was supported by a grant from the National Institutes of Health (GM45336) and core facilities by a National Cancer Institute Core grant (CA045508). We thank A. Chabes for the antibodies against Rnr4 and Sml1, H. Araki and J. Diffley for yeast strains, Cold Spring Harbor Laboratory DNA Sequencing Next Gen shared resource for high throughput sequencing, Microscopy, Flow Cytometry shared resources and Bioinformatics core for initial data analysis, Patty Wendel and James Building Staff for general assistance.

### DISCLOSURE DECLARATION (including any conflicts of interest)

No conflicts of interest exist

---

**Figure S1.**
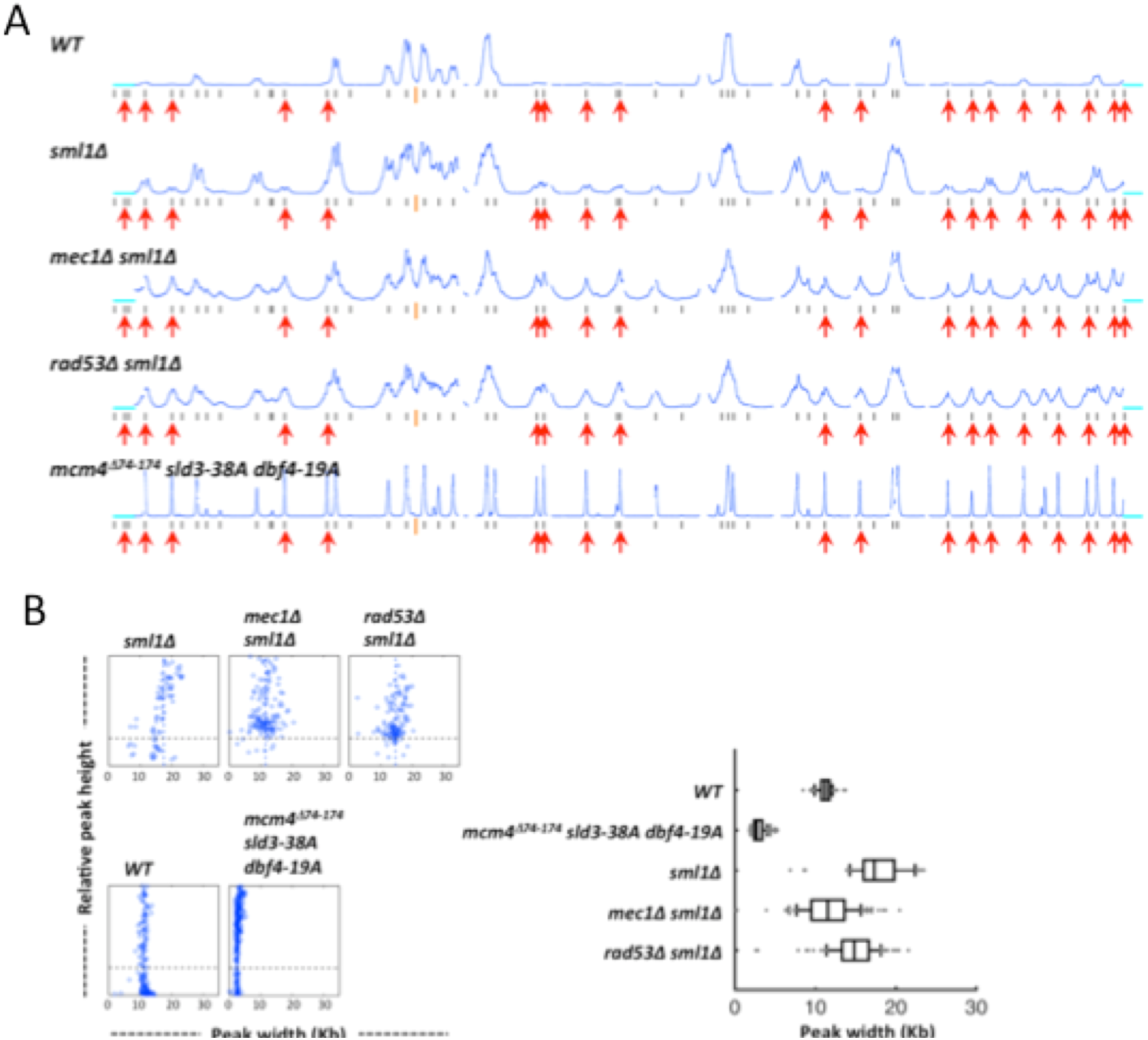
Late origin firing in HU in the absence of checkpoint signaling kinase. Yeast cells were synchronized in G1 phase and released in to YPD containing 0.2 M HU and 0.5 mM EdU for 90 min. (A) Replication profiles of chromosome IV for *WT*, *sml1Δ*, *mec1Δ sml1Δ*, *rad53Δ sml1Δ* and the *mcm4*^*Δ74-174*^ *sld3-38A dbf4-19A* triple mutant cells. The cyan lines mark the 30 kb region at the end of the chromosome that was masked. See main text for detail. (B) Distribution of fork progression from origins shown as individual width-height plots and box graph, excluding peaks with heights smaller than 30 % of the maximal height scale. The analysis was done using data with 30 kb at each end of each chromosome masked.

**Supplementary table 1.**

| Profiles in relevant genotype |  | Condition | mapped reads | smooth window | number peaks before filter | number peaks after filter | mean peak width after filter (kb) | SD (kb) | notes |
| --- | --- | --- | --- | --- | --- | --- | --- | --- | --- |
| Fig. 2A | WT | HU, 90 min | 13,311,114 | 1000 bp | 187 | 91 | 7.1 | 1.5 |  |
| Fig. 2A | *mcm4<sup>Δ74-174</sup>* | HU, 90 min | 7,289,123 | 1000 bp | 232 | 131 | 5.6 | 1.2 |  |
| Fig. 2A | *sld3-38A* | HU, 90 min | 10,503,231 | 1000 bp | 273 | 187 | 3.2 | 1 |  |
| Fig. 2A | *mcm4<sup>Δ74-174</sup> sld3-38A* | HU, 90 min | 10,875,046 | 1000 bp | 292 | 259 | 2.5 | 0.6 |  |
| Fig. 2A | *dbf4-19A* | HU, 90 min | 6,527,645 | 1000 bp | 221 | 113 | 5.9 | 2.8 |  |
| Fig. 2A | *mcm4<sup>Δ74-174</sup> dbf4-19A* | HU, 90 min | 11,190,863 | 1000 bp | 221 | 151 | 6.4 | 1.6 |  |
| Fig. 2A | *sld3-38A dbf4-19A* | HU, 90 min | 7,461,491 | 1000 bp | 299 | 239 | 2.2 | 0.9 |  |
| Fig. 2A | *mcm4<sup>Δ74-174</sup> sld3-38A dbf4-19A* | HU, 90 min | 8,734,875 | 1000 bp | 302 | 274 | 2.1 | 0.5 |  |
| Fig. 3A | WT | HU, 25 min | 4,612,750 | 500 bp | 271 | 149 | 1.3 | 0.2 |  |
| Fig. 3A | WT | HU, 50 min | 19,806,922 | 500 bp | 232 | 114 | 3.3 | 0.5 |  |
| Fig. 3A | WT | HU, 75 min | 26,924,268 | 500 bp | 184 | 92 | 6.8 | 1.1 |  |
| Fig. 3A | *mcm4<sup>Δ74-174</sup> sld3-38A dbf4-19A* | HU, 25 min | 6,182,622 | 500 bp | 302 | 218 | 1.1 | 0.2 |  |
| Fig. 3A | *mcm4<sup>Δ74-174</sup> sld3-38A dbf4-19A* | HU, 50 min | 17,944,322 | 500 bp | 305 | 265 | 1.1 | 0.3 |  |
| Fig. 3A | *mcm4<sup>Δ74-174</sup> sld3-38A dbf4-19A* | HU, 75 min | 26,404,584 | 500 bp | 292 | 265 | 2.1 | 0.6 |  |
| Fig. 3B | *sld3-38A dbf4-19A* | HU, 25 min | 15,199,022 | 500 bp | 259 | 119 | 1.5 | 0.4 |  |
| Fig. 3B | *sld3-38A dbf4-19A* | HU, 50 min | 18,232,289 | 500 bp | 295 | 187 | 1.5 | 0.7 |  |
| Fig. 3B | *sld3-38A dbf4-19A* | HU, 75 min | 14,651,214 | 500 bp | 293 | 252 | 2 | 1 |  |
| Fig. 3B | *mcm4<sup>Δ74-174</sup> sld3-38A dbf4-19A* | HU, 25 min | 17,966,386 | 500 bp | 295 | 215 | 1.2 | 0.3 |  |
| Fig. 3B | *mcm4<sup>Δ74-174</sup> sld3-38A dbf4-19A* | HU, 50 min | 22,584,515 | 500 bp | 305 | 263 | 1.2 | 0.4 |  |
| Fig. 3B | *mcm4<sup>Δ74-174</sup> sld3-38A dbf4-19A* | HU, 75 min | 17,510,882 | 500 bp | 291 | 265 | 1.9 | 0.6 |  |
| Fig. 4A | WT | HU, 90 min | 47,113,533 | 1000 bp | 168 | 62 | 8.6 | 0.9 |  |
| Fig. 4A | *mcm4<sup>Δ74-174</sup>* | HU, 90 min | 40,645,958 | 1000 bp | 201 | 73 | 7.1 | 0.8 |  |
| Fig. 4A | *sld3-38A* | HU, 90 min | 40,183,230 | 1000 bp | 271 | 128 | 3.9 | 0.9 |  |
| Fig. 4A | *mcm4<sup>Δ74-174</sup> sld3-38A* | HU, 90 min | 49,166,452 | 1000 bp | 286 | 211 | 2.8 | 0.6 |  |
| Fig. 4A | *mcm4<sup>Δ74-174, 4(SP→AP)</sup> sld3-38A* | HU, 90 min | 47,911,372 | 1000 bp | 194 | 80 | 7.5 | 1 |  |
| Fig. 4A | *mcm4<sup>Δ2-145</sup>* | HU, 90 min | 45,483,693 | 1000 bp | 62 | 15 | 20.4 | 3.9 |  |
| Fig. 4A | *mcm4<sup>Δ2-145</sup> sld3-38A* | HU, 90 min | 40,906,504 | 1000 bp | 166 | 60 | 9.9 | 1.4 |  |
| Fig. 5A | WT | HU, 90 min | 49,209,936 | 2000 bp | 161 | 73 | 11.2 | 0.9 |  |
| Fig. 5A | *mcm4<sup>Δ74-174</sup> sld3-38A dbf4-19A* | HU, 90 min | 36,950,160 | 1000 bp | 287 | 237 | 2.9 | 0.7 |  |
| Fig. 5A | *sml1Δ* | HU, 90 min | 41,858,795 | 2000 bp | 82 | 58 | 17.7 | 3.4 |  |
| Fig. 5A | *mec1Δ sml1Δ* | HU, 90 min | 41,429,448 | 2000 bp | 147 | 134 | 11.3 | 3.2 |  |
| Fig. 5A | *rad53Δ sml1Δ* | HU, 90 min | 41,234,875 | 2000 bp | 124 | 107 | 14.6 | 3.3 |  |
| Fig. S1A | WT | HU, 90 min | 49,209,936 | 2000 bp | 153 | 71 | 11.1 | 0.9 | ends masked |
| Fig. S1A | *mcm4<sup>Δ74-174</sup> sld3-38A dbf4-19A* | HU, 90 min | 36,950,160 | 1000 bp | 267 | 221 | 2.9 | 0.7 | ends masked |
| Fig. S1A | *sml1Δ* | HU, 90 min | 41,858,795 | 2000 bp | 79 | 57 | 17.6 | 3.4 | ends masked |
| Fig. S1A | *mec1Δ sml1Δ* | HU, 90 min | 41,429,448 | 2000 bp | 141 | 133 | 11.5 | 3.2 | ends masked |
| Fig. S1A | *rad53Δ sml1Δ* | HU, 90 min | 41,234,875 | 2000 bp | 120 | 107 | 14.7 | 3.1 | ends masked |
| Fig. 6A | WT | YPD, 25 min | 21,381,611 | 1000 bp | 206 | 99 | 7.2 | 1.2 |  |
| Fig. 6A | *mcm4<sup>Δ2-145</sup>* | YPD, 25 min | 13,296,061 | 1000 bp | 186 | 72 | 8.1 | 1.3 |  |
| Fig. 6A | *mcm4<sup>Δ2-174</sup>* | YPD, 25 min | 8,314,459 | 1000 bp | 207 | 136 | 7 | 1.3 |  |
| Fig. 6A | *mcm4<sup>Δ74-174</sup>* | YPD, 25 min | 25,590,224 | 1000 bp | 210 | 120 | 7.1 | 1 |  |
| Fig. 6A | *mcm4<sup>Δ74-174, 4(SP→AP)</sup>* | YPD, 25 min | 13,003,604 | 1000 bp | 174 | 76 | 8.8 | 1.5 |  |
| Fig. 6A | *mcm4<sup>Δ74-174, 4(SP→DP)</sup>* | YPD, 25 min | 21,982,699 | 1000 bp | 197 | 107 | 8.1 | 1.3 |  |
| Fig. 7A | WT | YPD, 25 min | 5,880,287 | 1000 bp | 236 | 98 | 4.8 | 1 |  |
| Fig. 7A | *mcm4<sup>Δ74-174</sup>* | YPD, 25 min | 8,532,204 | 1000 bp | 252 | 110 | 3.8 | 0.7 |  |
| Fig. 7A | *sld3-38A* | YPD, 25 min | 7,370,826 | 1000 bp | 237 | 97 | 4.7 | 1 |  |
| Fig. 7A | *mcm4<sup>Δ74-174</sup> sld3-38A* | YPD, 25 min | 3,769,402 | 1000 bp | 245 | 124 | 4.6 | 0.9 |  |
| Fig. 7A | *dbf4-19A* | YPD, 25 min | 17,717,050 | 1000 bp | 178 | 70 | 7.8 | 1.9 |  |
| Fig. 7A | *mcm4<sup>Δ74-174</sup> dbf4-19A* | YPD, 25 min | 6,040,131 | 1000 bp | 231 | 114 | 5.3 | 1.2 |  |
| Fig. 7A | *sld3-38A dbf4-19A* | YPD, 25 min | 4,409,073 | 1000 bp | 216 | 73 | 4.7 | 1 |  |
| Fig. 7A | *mcm4<sup>Δ74-174</sup> sld3-38A dbf4-19A* | YPD, 25 min | 10,194,071 | 1000 bp | 239 | 100 | 4.4 | 0.8 |  |
| Fig. 8A | WT | MMS, 50 min | 17,523,516 | 2000 bp | 82 | 51 | 17.3 | 1.7 |  |
| Fig. 8A | *mcm4<sup>Δ2-145</sup>* | MMS, 50 min | 17,782,954 | 2000 bp | 17 | 10 | 29.4 | 12.4 |  |
| Fig. 8A | *mcm4<sup>Δ2-174</sup>* | MMS, 50 min | 21,104,155 | 2000 bp | 17 | 15 | 22.1 | 13 |  |
| Fig. 8A | *mcm4<sup>Δ74-174</sup>* | MMS, 50 min | 19,871,276 | 2000 bp | 124 | 74 | 14.2 | 2.3 |  |
| Fig. 8A | *mcm4<sup>Δ74-174, 4(SP→AP)</sup>* | MMS, 50 min | 15,214,419 | 2000 bp | 25 | 13 | 27.6 | 8.8 |  |
| Fig. 8A | *mcm4<sup>Δ74-174, 4(SP→DP)</sup>* | MMS, 50 min | 20,407,279 | 2000 bp | 63 | 42 | 20 | 3.1 |  |
| Fig. 9A | WT | MMS, 50 min | 25,602,878 | 2000 bp | 78 | 47 | 16.9 | 2.4 |  |
| Fig. 9A | *mcm4<sup>Δ74-174</sup>* | MMS, 50 min | 19,140,949 | 2000 bp | 117 | 62 | 13.7 | 2.1 |  |
| Fig. 9A | *sld3-38A* | MMS, 50 min | 24,865,918 | 2000 bp | 159 | 102 | 11.6 | 2.2 |  |
| Fig. 9A | *mcm4<sup>Δ74-174</sup> sld3-38A* | MMS, 50 min | 26,546,733 | 2000 bp | 190 | 146 | 9.3 | 1.4 |  |
| Fig. 9A | *dbf4-19A* | MMS, 50 min | 29,456,510 | 2000 bp | 49 | 24 | 24 | 4.5 |  |
| Fig. 9A | *mcm4<sup>Δ74-174</sup> dbf4-19A* | MMS, 50 min | 27,231,721 | 2000 bp | 90 | 55 | 17.3 | 4.3 |  |
| Fig. 9A | *sld3-38A dbf4-19A* | MMS, 50 min | 20,216,769 | 2000 bp | 176 | 117 | 10.9 | 3.9 |  |
| Fig. 9A | *mcm4<sup>Δ74-174</sup> sld3-38A dbf4-19A* | MMS, 50 min | 31,625,044 | 2000 bp | 199 | 177 | 9 | 1.7 |  |

**Supplementary table 2.**
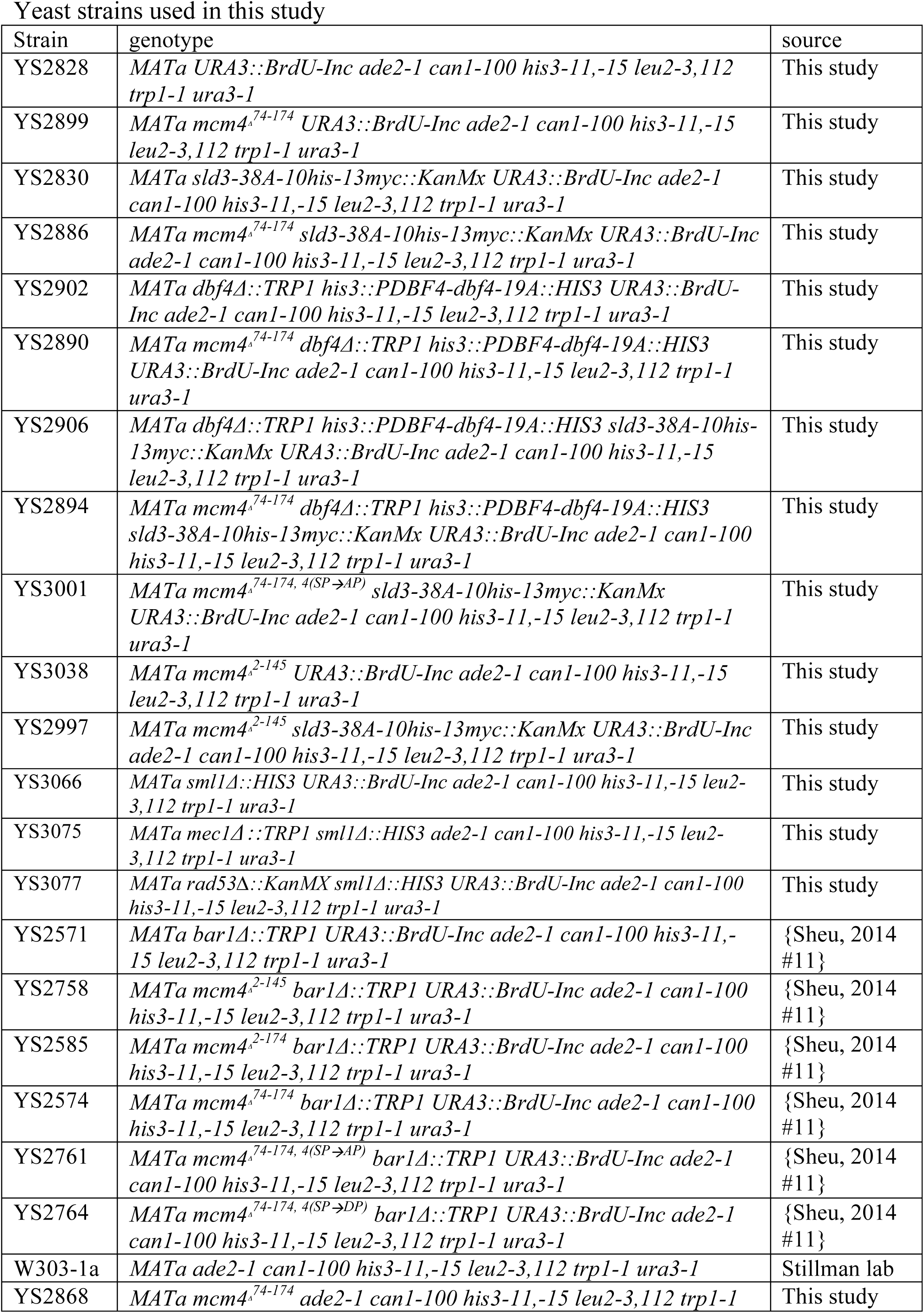
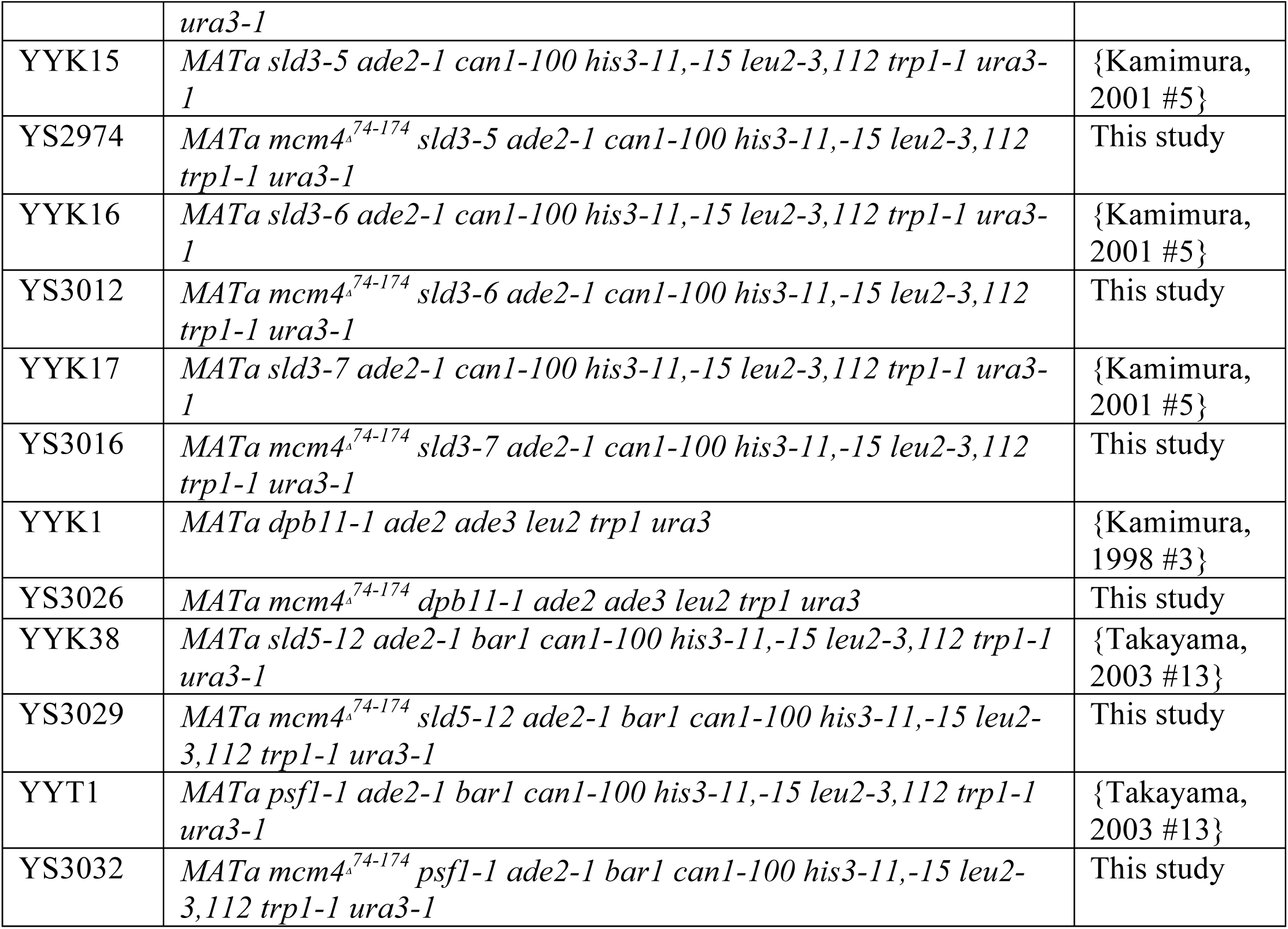
Yeast strains used in this study

## REFERENCES

Alvino GM, Collingwood D, Murphy JM, Delrow J, Brewer BJ, Raghuraman MK. 2007. Replication in hydroxyurea: it’s a matter of time. Mol Cell Biol 27: 6396–6406.

Chabes A, Stillman B. 2007. Constitutively high dNTP concentration inhibits cell cycle progression and the DNA damage checkpoint in yeast Saccharomyces cerevisiae. Proc Natl Acad Sci U S A 104: 1183–1188.

Devault A, Gueydon E, Schwob E. 2008. Interplay between S-cyclin-dependent kinase and Dbf4-dependent kinase in controlling DNA replication through phosphorylation of yeast Mcm4 N-terminal domain. Mol Biol Cell 19: 2267–2277.

Diffley JF. 2011. Quality control in the initiation of eukaryotic DNA replication. Philos Trans R Soc Lond B Biol Sci 366: 3545–3553.

Dohrmann PR, Oshiro G, Tecklenburg M, Sclafani RA. 1999. RAD53 regulates DBF4 independently of checkpoint function in Saccharomyces cerevisiae. Genetics 151: 965–977.

Donaldson AD, Raghuraman MK, Friedman KL, Cross FR, Brewer BJ, Fangman WL. 1998. CLB5-dependent activation of late replication origins in S. cerevisiae. Mol Cell 2: 173–182.

Heller RC, Kang S, Lam WM, Chen S, Chan CS, Bell SP. 2011. Eukaryotic origin-dependent DNA replication in vitro reveals sequential action of DDK and S-CDK kinases. Cell 146: 80–91.

Jones DR, Prasad AA, Chan PK, Duncker BP. 2010. The Dbf4 motif C zinc finger promotes DNA replication and mediates resistance to genotoxic stress. Cell Cycle 9: 2018–2026.

Kamimura Y, Masumoto H, Sugino A, Araki H. 1998. Sld2, which interacts with Dpb11 in Saccharomyces cerevisiae, is required for chromosomal DNA replication. Mol Cell Biol 18: 6102–6109.

Kamimura Y, Tak YS, Sugino A, Araki H. 2001. Sld3, which interacts with Cdc45 (Sld4), functions for chromosomal DNA replication in Saccharomyces cerevisiae. EMBO J 20: 2097–2107.

Kanemaki M, Labib K. 2006. Distinct roles for Sld3 and GINS during establishment and progression of eukaryotic DNA replication forks. EMBO J 25: 1753–1763.

Kihara M, Nakai W, Asano S, Suzuki A, Kitada K, Kawasaki Y, Johnston LH, Sugino A. 2000. Characterization of the yeast Cdc7p/Dbf4p complex purified from insect cells. Its protein kinase activity is regulated by Rad53p. J Biol Chem 275: 35051–35062.

Lopez-Mosqueda J, Maas NL, Jonsson ZO, Defazio-Eli LG, Wohlschlegel J, Toczyski DP. 2010. Damage-induced phosphorylation of Sld3 is important to block late origin firing. Nature 467: 479–483.

Mantiero D, Mackenzie A, Donaldson A, Zegerman P. 2011. Limiting replication initiation factors execute the temporal programme of origin firing in budding yeast. EMBO J 30: 4805–4814.

Masai H, Taniyama C, Ogino K, Matsui E, Kakusho N, Matsumoto S, Kim JM, Ishii A, Tanaka T, Kobayashi T et al. 2006. Phosphorylation of MCM4 by Cdc7 kinase facilitates its interaction with Cdc45 on the chromatin. J Biol Chem 281: 39249–39261.

Muramatsu S, Hirai K, Tak YS, Kamimura Y, Araki H. 2010. CDK-dependent complex formation between replication proteins Dpb11, Sld2, Pol (epsilon), and GINS in budding yeast. Genes Dev 24: 602–612.

Poli J, Tsaponina O, Crabbe L, Keszthelyi A, Pantesco V, Chabes A, Lengronne A, Pasero P. 2012. dNTP pools determine fork progression and origin usage under replication stress. EMBO J 31: 883–894.

Randell JC, Fan A, Chan C, Francis LI, Heller RC, Galani K, Bell SP. 2010. Mec1 is one of multiple kinases that prime the Mcm2-7 helicase for phosphorylation by Cdc7. Mol Cell 40: 353–363.

Rhind N, Gilbert DM. 2013. DNA replication timing. Cold Spring Harb Perspect Biol 5: a010132.

Sanchez Y, Desany BA, Jones WJ, Liu Q, Wang B, Elledge SJ. 1996. Regulation of RAD53 by the ATM-like kinases MEC1 and TEL1 in yeast cell cycle checkpoint pathways. Science 271: 357–360.

Santocanale C, Diffley JF. 1998. A Mec1- and Rad53-dependent checkpoint controls late-firing origins of DNA replication. Nature 395: 615–618.

Sheu YJ, Kinney JB, Lengronne A, Pasero P, Stillman B. 2014. Domain within the helicase subunit Mcm4 integrates multiple kinase signals to control DNA replication initiation and fork progression. Proc Natl Acad Sci U S A 111: E1899–1908.

Sheu YJ, Stillman B. 2006. Cdc7-Dbf4 phosphorylates MCM proteins via a docking site-mediated mechanism to promote S phase progression. Mol Cell 24: 101–113.

Sheu YJ, Stillman B. 2010. The Dbf4-Cdc7 kinase promotes S phase by alleviating an inhibitory activity in Mcm 4. Nature 463: 113–117.

Siow CC, Nieduszynska SR, Muller CA, Nieduszynski CA. 2012. OriDB, the DNA replication origin database updated and extended. Nucleic Acids Res 40: D682–686.

Takayama Y, Kamimura Y, Okawa M, Muramatsu S, Sugino A, Araki H. 2003. GINS, a novel multiprotein complex required for chromosomal DNA replication in budding yeast. Genes Dev 17: 1153–1165.

Tanaka S, Araki H. 2013. Helicase activation and establishment of replication forks at chromosomal origins of replication. Cold Spring Harb Perspect Biol 5: a010371.

Tanaka S, Nakato R, Katou Y, Shirahige K, Araki H. 2011. Origin association of Sld3, Sld7, and Cdc45 proteins is a key step for determination of origin-firing timing. Curr Biol 21: 2055–2063.

Tanaka S, Umemori T, Hirai K, Muramatsu S, Kamimura Y, Araki H. 2007. CDK-dependent phosphorylation of Sld2 and Sld3 initiates DNA replication in budding yeast. Nature 445: 328–332.

Tercero JA, Diffley JF. 2001. Regulation of DNA replication fork progression through damaged DNA by the Mec1/Rad53 checkpoint. Nature 412: 553–557.

Tercero JA, Longhese MP, Diffley JF. 2003. A central role for DNA replication forks in checkpoint activation and response. Mol Cell 11: 1323–1336.

Varrin AE, Prasad AA, Scholz RP, Ramer MD, Duncker BP. 2005. A mutation in Dbf4 motif M impairs interactions with DNA replication factors and confers increased resistance to genotoxic agents. Mol Cell Biol 25: 7494–7504.

Viggiani CJ, Aparicio OM. 2006. New vectors for simplified construction of BrdU-Incorporating strains of Saccharomyces cerevisiae. Yeast 23: 1045–1051.

Weinreich M, Stillman B. 1999. Cdc7p-Dbf4p kinase binds to chromatin during S phase and is regulated by both the APC and the RAD53 checkpoint pathway. EMBO J 18: 5334–5346.

Zegerman P, Diffley JF. 2007. Phosphorylation of Sld2 and Sld3 by cyclin-dependent kinases promotes DNA replication in budding yeast. Nature 445: 281–285.

Zegerman P, Diffley JF. 2009. DNA replication as a target of the DNA damage checkpoint. DNA Repair (Amst) 8: 1077–1088.

Zegerman P, Diffley JF. 2010. Checkpoint-dependent inhibition of DNA replication initiation by Sld3 and Dbf4 phosphorylation. Nature 467: 474–478.

Zhao X, Muller EG, Rothstein R. 1998. A suppressor of two essential checkpoint genes identifies a novel protein that negatively affects dNTP pools. Mol Cell 2: 329–340.

Zhong Y, Nellimoottil T, Peace JM, Knott SR, Villwock SK, Yee JM, Jancuska JM, Rege S, Tecklenburg M, Sclafani RA et al. 2013. The level of origin firing inversely affects the rate of replication fork progression. J Cell Biol 201: 373–383.

